# Direct Quantification of *in vivo* Mutagenesis and Carcinogenesis Using Duplex Sequencing

**DOI:** 10.1101/2020.06.28.176685

**Authors:** Charles C. Valentine, Robert R. Young, Mark R. Fielden, Rohan Kulkarni, Lindsey N. Williams, Tan Li, Sheroy Minocherhomji, Jesse J. Salk

## Abstract

The ability to accurately measure mutations is critical for basic research and identification of potential drug and chemical carcinogens. Current methods for *in vivo* quantification of mutagenesis are limited because they rely on transgenic rodent systems that are low-throughput, expensive, prolonged, and don’t fully represent other species such as humans. Next generation sequencing (NGS) is a conceptually attractive alternative for mutation detection in the DNA of any organism, however, the limit of resolution for standard NGS is poor. Technical error rates (~1×10^−3^) of NGS obscure the true abundance of somatic mutations, which can exist at per-nucleotide frequencies ≤1×10^−7^. Using Duplex Sequencing, an extremely accurate error-corrected NGS (ecNGS) technology, we were able to detect mutations induced by 3 carcinogens in 5 tissues of 2 strains of mice within 31 days following exposure. We observed a strong correlation between mutation induction measured by Duplex Sequencing and the gold-standard transgenic rodent mutation assay. We identified exposure-specific mutation spectra of each compound through trinucleotide patterns of base substitution. We observed variation in mutation susceptibility by genomic region, as well as by DNA strand. We also identified the primordial signs of carcinogenesis in a cancer-predisposed strain of mice, as evidenced by clonal expansions of cells carrying an activated oncogene, less than a month after carcinogen exposure. These findings demonstrate that ecNGS is a powerful method for sensitively detecting and characterizing mutagenesis and the early clonal evolutionary hallmarks of carcinogenesis. Duplex Sequencing can be broadly applied to chemical safety testing, basic mutational research, and related clinical uses.

**SIGNIFICANCE STATEMENT:** Error-corrected next generation sequencing (ecNGS) can be used to rapidly detect and quantify the *in vivo* mutagenic impact of environmental exposures or endogenous processes in any tissue, from any species, at any genomic location. The greater speed, higher scalability, richer data outputs, as well as cross-species and cross-locus applicability of ecNGS compared to existing methods make it a powerful new tool for mutational research, regulatory safety testing, and emerging clinical applications.

## INTRODUCTION

Carcinogenesis is rooted in somatic evolution. Cell populations bearing stochastically-arising genetic mutations undergo iterative waves of natural selection that enrich for mutants which confer a phenotype of preferential survival or proliferation^1^. The probability of cancer can be increased by carcinogens—exogenous exposures that either increase the abundance of mutations or facilitate a cell’s ability to proliferate upon selective pressures. Many chemicals induce DNA damage, thereby increasing the rate of potentially oncogenic DNA replication errors^2^. The same is true for many forms of radiation^3^. Non-mutagenic/non-genotoxic carcinogens act through a variety of secondary mechanisms such as inhibition of the immune system, cell cycle overdrive to bypass normal DNA replication checkpoints, and induction of inflammation which may lead to both increased cellular proliferation and DNA damage, among others^4^

Preclinical genotoxicity and carcinogenicity testing of new compounds is often required before regulatory authority approval and subsequent human exposure^5,6^. However, this standard is slow and expensive; even in rodents, it takes years to reach the endpoint of tumor formation. Over the past fifty years, a variety of approaches have been developed to more quickly assess biomarkers of genotoxicity or cancer risk by assaying DNA reactivity or mutagenic potential as surrogate endpoints for regulatory decision-making^7^ The most rapid and inexpensive of such methods include *in vitro* bacterial-based mutagenesis assays *(e.g.* the Ames test). Other *in vitro* and *in vivo* assays for mutation, chromosomal aberration induction, strand breakage, and formation of micronuclei are also available, however, their sensitivity and specificity for predicting human cancer risk is only modest.

*In vivo,* internationally accepted^5^ mutagenesis assays using transgenic rodents (TGR) provide a powerful approximation of oncogenic risk, as they reflect whole-organism biology, but are also highly complex test systems^8^. Transgenic rodent mutagenesis assays require maintenance of multiple generations of animals bearing an artificial reporter gene, animal exposure to the test compound, necropsy several weeks after exposure, isolation of the integrated genetic reporter by phage packaging, and transfection of the phage into *E. coli* for plaque-counting on many petri dishes under permissive and non-permissive selection conditions to finally obtain a mutant frequency readout. Although effective, the infrastructure and expertise required for managing a protocol which carries host DNA through three different kingdoms of life has not been amenable to ubiquitous adoption.

Directly measuring ultra-rare somatic mutations from extracted DNA while not being restricted by genomic locus, tissue, or organism *(i.e.* could be equally applied to rodents or humans) is appealing yet is currently impossible with conventional next-generation DNA sequencing (NGS). Standard NGS has a technical error rate (~1×10^−3^) well above the true per-nucleotide mutant frequency of normal tissues (<1×10^−7^)^9^. New technologies for error-corrected next generation sequencing (ecNGS) have shown great promise for low frequency mutation detection in fields such as oncology and, conceptually, could be applied to genetic toxicology^10,11^. Duplex Sequencing (DS), in particular, is an error correction method that achieves extremely high accuracy by comparing reads derived from both original strands of DNA molecules to produce duplex consensus sequences that better represent the true sequence of the source DNA molecule. DS achieves a sensitivity and specificity several orders of magnitude greater than other methods that do not leverage paired-strand information; it is uniquely able to resolve mutants at the real-world frequencies produced by mutagens: on the order of one-in-ten-million^12^.

In this study we tested the feasibility of using Duplex Sequencing to measure the effect of genotoxicants *in vivo.* We assessed the DNA of two strains of mice which were treated with three different mutagenic carcinogens, each with distinct modes of action, and examined five different tissue types, to generate a total of almost ten-billion error-corrected nucleotides worth of data. In addition to comparing mutant frequencies with those obtained from a gold-standard TGR assay, we explored data types not possible with traditional assays, including mutant spectra, trinucleotide signatures, and variations in the relative mutagenic sensitivity around the genome. Our findings illustrate the richness of genotoxicity data that can be obtained directly from genomic DNA. Finally, we highlight an opportunity in applying ecNGS to drug and chemical safety testing, as well as broader applications in the research or clinical settings, for the detection of phenomena related to both mutagenesis and carcinogenesis.

## RESULTS

### Experimental overview

Current *in vivo* transgenic rodent (TGR) mutagenesis detection assays measure the potential of a test article to generate mutations in a selectable reporter gene. Traditional two-year carcinogenicity studies measure the ability of an agent to induce gross tumors in mice and rats. We designed two parallel mouse cohort studies to assess whether Duplex Sequencing (DS) of genomic DNA could be used as an alternate method of quantifying both mutagenesis and early tumor-precursor formation (**Fig. 1**). We selected two transgenic strains of mice: Big Blue^®^, a C56BL/6-derived strain bearing approximately 40 integrated copies of lambda phage per cell and Tg-rasH2, a cancer-predisposed strain carrying approximately 4 copies of the human *HRAS* proto-oncogene^13,14^. The Big Blue mouse is one of the three most frequently used strains in the TGR mutagenesis assay and the Tg-rasH2 mouse is used for accelerated 6-month carcinogenicity studies. Animals were dosed for up to twenty-eight days with one of three mutagenic chemicals (or vehicle control) and then were sacrificed and necropsied. Genomic DNA was isolated from various frozen tissues for subsequent mutational analysis (**Table 1**). The rationale for the selection of the specific strains, chemicals, tissues, and genic targets to be sequenced is detailed in the sections that follow.

**Figure 1.**
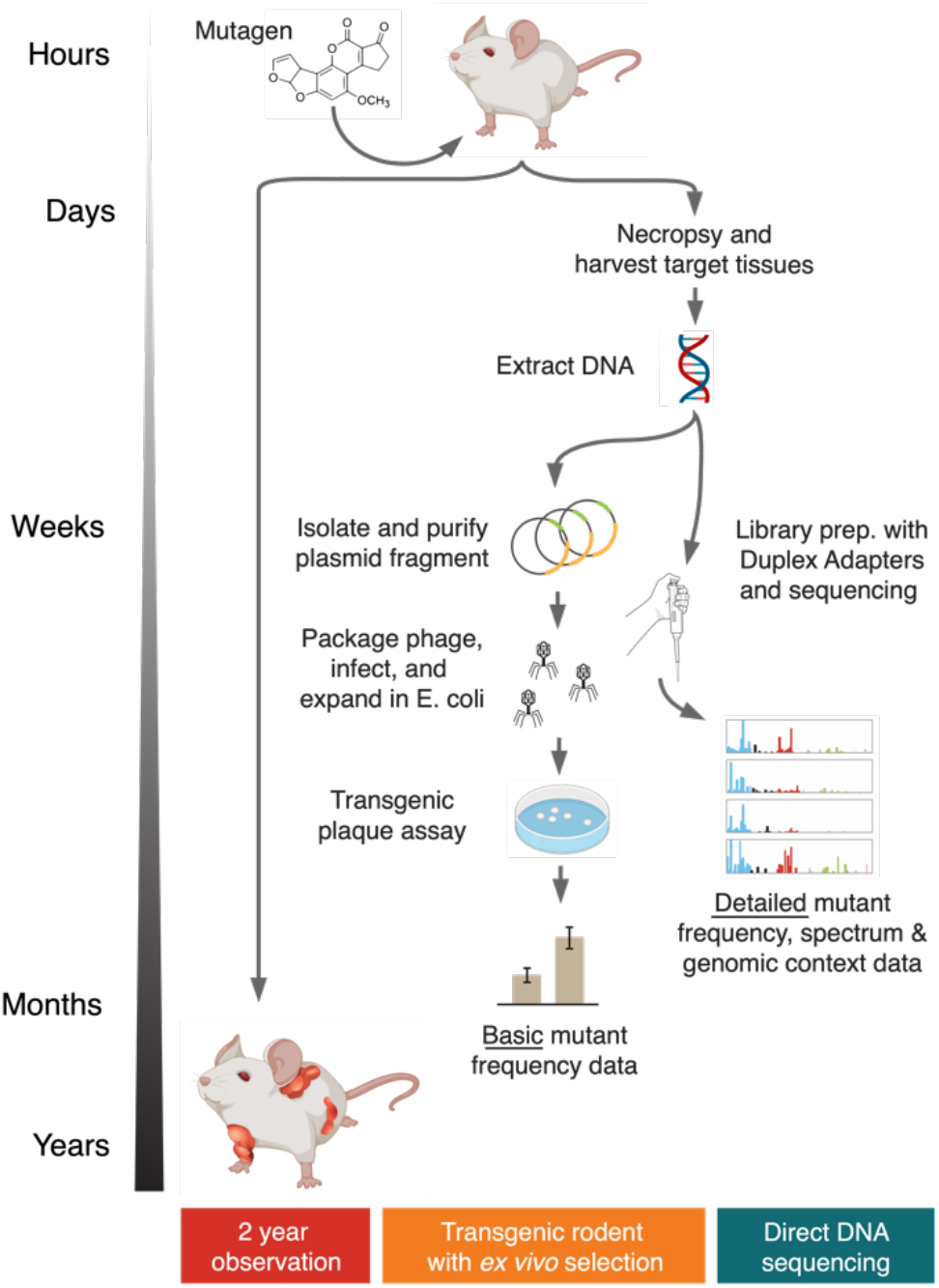
Approaches for *in vivo* carcinogenicity and mutagenicity testing. The gold-standard for chemical cancer risk assessment involves exposing rodents to a test compound and assessing an increase in tumors relative to controls. While effective, the approach takes two years. Mutagenicity is one accepted surrogate for carcinogenicity risk which is more rapidly assessable. At present, transgenic rodent assays are the most validated *in vivo* mutagenesis measurement approach. A genetically modified strain carrying an artificial reporter gene is mutagen-exposed, necropsied, and the reporter is recovered from extracted DNA. Reporter DNA is then packaged into phage and transfected into bacteria for readout by counting total and mutation-bearing plaques following incubation at mutant-selectable temperature conditions. A simple mutant frequency can be determined within months, but the assay is complex. An ideal *in vivo* mutagenesis assay would be DNA sequencing-based and able to directly measure mutation abundance and spectrum in any tissue of any species without the need for genetic engineering or selection.

**Table 1.** Summary of all samples along with cohort-level metadata.

|  | Big Blue | Tg-rasH2 | Total |
| --- | --- | --- | --- |
| Tissues<br>(samples per group) | Liver (15)<br>Marrow (17) | Lung (10)<br>Spleen (10)<br>Blood (10) | 5 tissue types |
| Treatment<br>(samples per group) | B[$\alpha$]P (10)<br>ENU (11)<br>VC (11) | Urethane (15)<br>VC (15) | 3 mutagens |
| Number Samples | 32 | 30 | 62 samples |
| Endogenous Targets | *Ctnnb1*<br>*Hp*<br>*Polr1c*<br>*Rho* | *Ctnnb1*<br>*Hp*<br>*Hras*<br>*Kras*<br>*Nras*<br>*Polr1c*<br>*Rho* | 7 endogenous targets |
| Transgenic Targets | Lambda<br>bacteriophage<br>*cII* | Human *HRAS* | 2 transgenic targets |
| Duplex Base Pairs | 4,716,990,836 | 4,923,565,684 | 9,640,556,520 |

### Duplex Sequencing yields comparable results to the transgenic rodent assay

We compared the frequencies of chemically induced mutations measured by a conventional plaque-based TGR assay (Big Blue) against those obtained by Duplex Sequencing of the Big Blue reporter gene *(cII)* after isolation from mouse gDNA in the absence of any *in vitro* selection.

Eighteen Big Blue mice were treated with either vehicle control (VC, olive oil), benzo[α]pyrene (B[α]P) or N-ethyl-N-nitrosourea (ENU) for up to 28 days (see **Materials and Methods**). We selected B[α]P and ENU based on their historical use as positive controls in many early studies of mutagenesis^15^ and because they are recommended by OECD TG 488 for demonstration of proficiency at detecting *in vivo* mutagenesis with TGR assays^5^. We evaluated bone marrow and liver tissue. The former was selected based on its high cell division rate and the latter based its slower cell division rate and the presence of enzymes necessary for converting some non-reactive mutagenic compounds into their DNA-reactive metabolites. Corresponding plaque-based *cII* gene mutant frequency data using the Big Blue TGR plaque-based assay was collected on all samples (**Supplemental Table 1**).

Genomic DNA was ultrasonically sheared and processed using a previously reported DS approach^16^, which included enrichment for genic targets via hybrid capture (see **Materials and Methods**). All samples were initially investigator-blinded with regard to treatment group. In this first experiment, we sequenced the multi-copy *cII* transgene to a mean duplex depth (*i.e.* single duplex source molecule, deduplicated, coverage) of 39,668x per sample. Duplex Sequencing mutant frequency per sample was calculated as the total number of unique non-reference nucleotides detected among all duplex reads of the *cII* gene divided by the total number of duplex base pairs of the *cII* gene sequenced.

The mean per-nucleotide mutant frequency measured by DS in the VC, B[α]P, and ENU-exposed groups was 1.48×10^−7^, 1.16×10^−6^ (7.84-fold increase over VC) and 1.27×10^−6^ (8.58-fold increase over VC) respectively. The mean fold-increase detected between vehicle control and mutagen-exposed groups was similar to that measured by the conventional plaque assay, with per-gene mutant frequencies for VC, B[α]P, and ENU averaging 4.09×10^−5^, 4.42×10^−4^ (10.81 fold-increase over VC), and 3.06×10^−4^ (7.48-fold increase over VC), respectively (**Fig. 2A**). The extent of induction by both assays was dependent on the tissue type. Bone marrow cells, with their higher proliferation rate, accumulated mutations at 3.75 and 2.48 times the rate of the slower-dividing cells from the liver for B[α]P and ENU, respectively.

**Figure 2.**
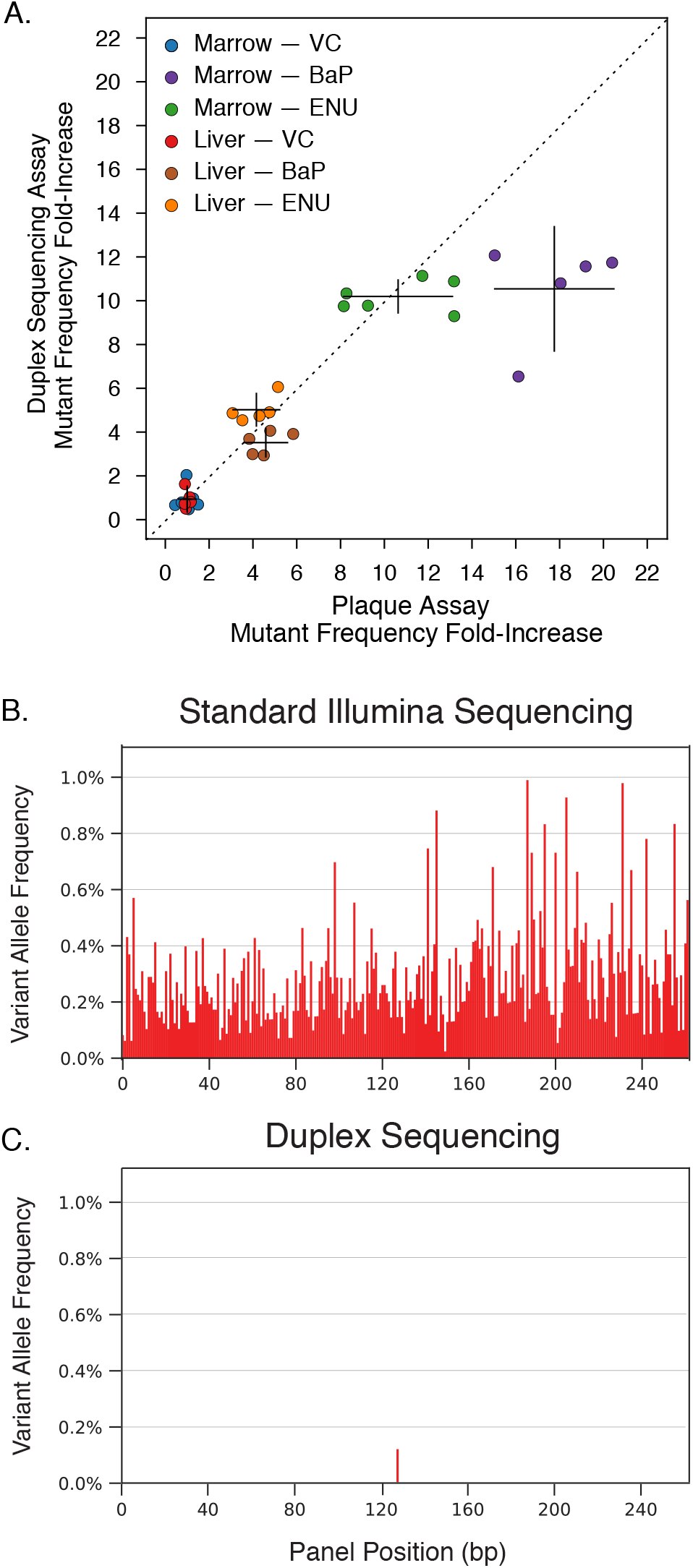
Comparison of Duplex Sequencing and the transgenic rodent assay for quantifying *in vivo* chemical mutagenesis. **(A)** Duplex Sequencing (DS) of a transgenic reporter gene *(cII)* relied upon by the Big Blue mouse TGR assay yields a similar fold-induction of mutations in response to chemical mutagenesis as the readouts from the plaque-based assay. Error bars reflect 95% confidence interval. **(B)** Standard DNA sequencing has an error rate of between 0.1% and 1% which obscures the presence of genuine low frequency mutations. Shown is conventional sequencing data from a representative 250 base pair section of the human *HRAS* transgene from the lung of a Tg-rasH2 mouse in the present study. Each bar corresponds to a nucleotide position. The height of each bar corresponds to the allele fraction of non-reference bases at that position when sequenced to >100,000x depth. Every position appears to be mutated at some frequency; nearly all of these are errors. **(C)** When the same sample is processed with DS, only a single authentic mutation remains.

The extent of correlation between the fold change mutation induction of the two methods (R^2^ = 0.898) was encouraging given that the assays measure mutant frequency via two fundamentally different approaches. Duplex Sequencing genotypes millions of unique nucleotides to assess the proportion that are mutated whereas the plaque assay measures the proportion of phage-packaged *cII* genes that bear at least one mutation that sufficiently disrupts the function of the cII protein to result in phenotypic plaque formation. Put another way, mutations that are disruptive enough to prevent packaging or phage expression in *E. coli*, or those that are synonymous or otherwise have no functional impact on the cII protein, will not be scored.

One difference observed between the two methods was an attenuation of response to B[α]P in the marrow group by DS. This might be explained by an artificial skew due to the fold-increase calculation used, whereby slight variations in the frequency of VC will have disproportionately large effects on fold-increase measures, but could also be wholly biological. It is also conceivable that DNA adducts, or sites of true *in vivo* mismatches could be artefactually “fixed” into doublestranded mutations when passaging reporter fragments through *E. coli* in the TGR assay and that this effect is amplified as overall mutant frequency increases. Duplex Sequencing, based in its fundamental error-correction principle, will not call adducted DNA bases as mutations when directly sequencing the *cII* genomic DNA, since a mutation has not yet formed on both strands of the DNA molecule.

Nevertheless, the overall correlation between DS and TGR assays was high and the mutant frequency measured in the VC samples by DS, on the order of one-per-ten-million mutant nucleotides sequenced, was ten-thousand-fold below the average technical error rate of standard NGS (**Fig. 2B, C**). No difference in mutant frequency or spectrum between control and exposed samples could be detected when analyzing the data from either raw sequencing reads or ecNGS methods that do not account for complementary strand information (single-strand consensus sequencing) (**Fig. S1**).

### Duplex Sequencing detects similar base substitution spectra between gDNA and mutant plaques in the TGR assay

The types of base substitution changes that are induced is an important element of mutagenesis testing. What might appear as a lack of overall effect on mutant frequency can sometimes be clearly appreciated when analyzing a dataset for the individual frequencies of specific transitions and transversions. Mutation spectra can also provide mechanistic insight into the nature of a mutagen. Although laborious, it is possible with plaque assays to characterize mutation spectra by picking and sequencing the clonal phage populations of many individual plaques or plaque pools^17,18^. Because mutations in plaques have been functionally selected and the transgenic target is relatively small, it is possible that the spectral representation is skewed relative to a nonselection-based assay.

To assess whether mutation spectra are consistent between DS and TGR assays, we physically isolated, pooled, and sequenced (also with DS) 3,510 *cII* mutant plaques derived from Big Blue rodents exposed to VC, B[α]P, and ENU. We then compared the mutation spectra between the DS-analyzed mutant plaques and the DS-analyzed gDNA.

The base substitution spectra detected in the *cII* gene by both approaches were highly similar between methods (p-value >0.999, chi-squared test) (**Fig. S2**), and yielded patterns consistent with expectations based on prior literature for both B[α]P^19,20^, an agent with reactive metabolites that intercalate DNA, similar to Aflatoxin B_1_, and the alkylating agent ENU^21,22^. The majority of base substitutions observed following B[α]P exposure were characteristic G·C→T·A transversions (61.3% by DS, 57.0% by TGR), G·C→C·G transversions (17.5% by DS, 25.5% by TGR) and G·C→A·T transitions (16.2% by DS, 11.6% by TGR). The normally uncommon base substitutions with adenine or thymine as reference were increased in all ENU exposed samples. The canonical transition that identifies ENU mutagenesis, C·G→T·A, was present at 32.2% by DS and 27.0% by TGR. These data add further weight of evidence that the mutations identified by DS reflect authentic biology and not technical artifacts.

### Duplex Sequencing detects functional classes of cII mutants undetected by the plaque assay

The eponymously named TGR assays rely on a transgenic reporter cassette which can be recovered from genomic DNA. It is the ratio of mutant to wild-type genes, as inferred through phenotypically scoreable plaques, which permits the calculation of a mutant frequency^23–26^. While these systems readily identify a subset of mutations in the reporter, others will not disrupt protein function and remain undetectable. Given that the primary use of TGR assays has been for relative, rather than absolute, mutational comparison between exposed and unexposed animals, the non-functional subset of mutants has historically been considered irrelevant.

Yet with the increasing interest in more complex multi-nucleotide mutational spectra^27^, the functional scoring of every base becomes essential, given that a specific sequence may rarely, or never occur in a small reporter region. Duplex Sequencing does not have this limitation since there is no selection post-DNA extraction; all possible single nucleotide variants, multi-nucleotide variants, and indels can be equally well identified.

To illustrate the impact of TGR selection on mutant recovery, we plotted the functional class of all *cII* mutations identified by Duplex Sequencing of either genomic DNA obtained directly from mouse samples (**Fig. 3A**) or from a pool of 3,510 individual mutant plaques that were isolated post-selection (**Fig. 3B**). In the TGR plaque assay, the mutations were almost exclusively nonsense or missense across the entire 291 nucleotides of the *cII* gene (*i.e.* expected to result in the loss of *cII* protein function). Only a small number of synonymous base changes were identified, and these were always accompanied by a concomitant disruptive mutation elsewhere in the gene. Exceptionally few mutations were found at the N- and C-termini of the *cII* gene, presumably due to their lesser importance to protein function. In contrast, Duplex Sequencing detected mutations of all functional classes at the expected non-synonymous to synonymous (dN/dS) ratio along the entire length of the gene, including the termini regions.

**Figure 3.**
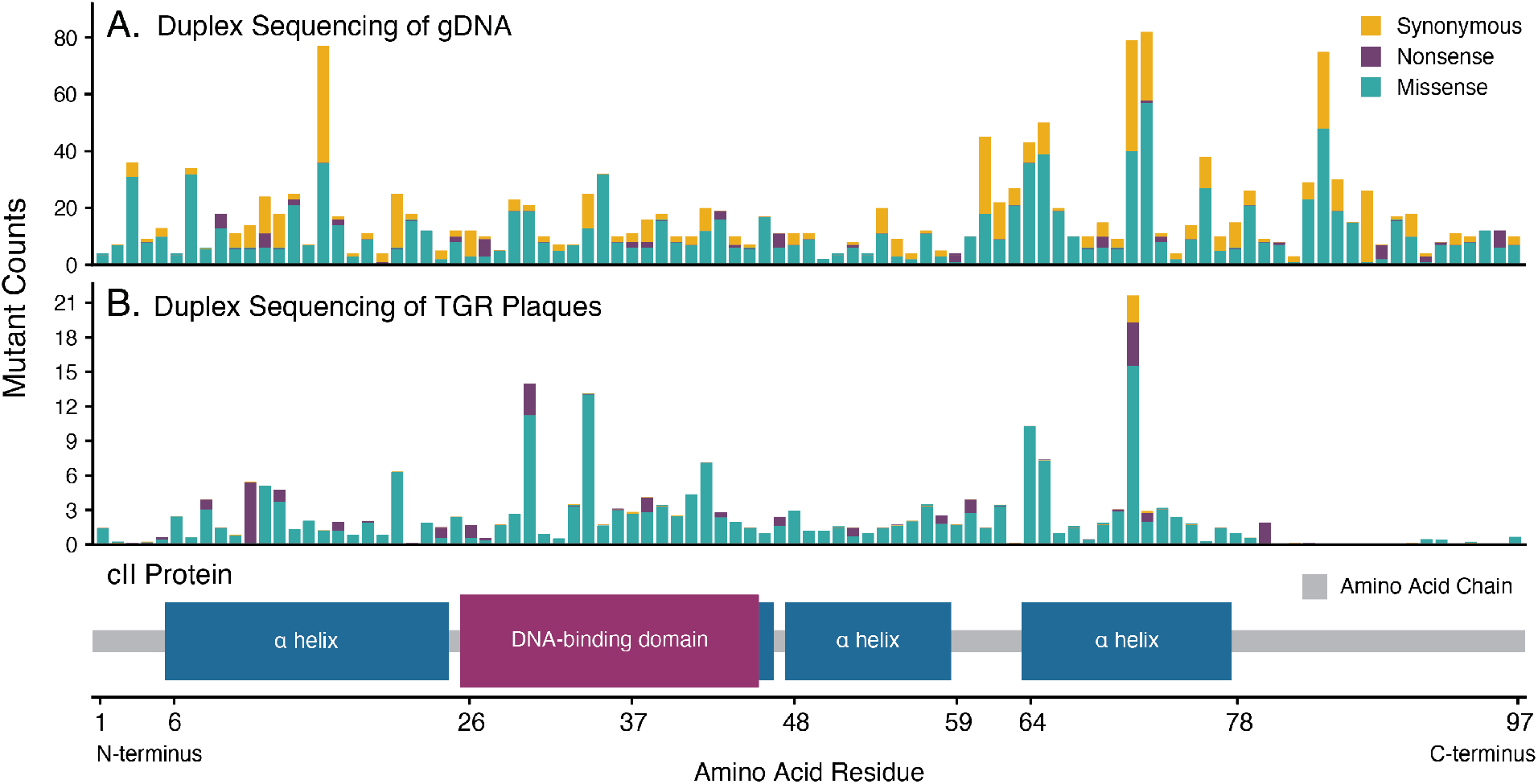
Duplex Sequencing is agnostic to reporter gene function whereas the TGR assay counts only phenotypically selectable mutations. **(A)** The distribution of all mutations identified by Duplex Sequencing of *cII* from genomic DNA across all Big Blue tissues and treatment groups is shown by codon position and functional consequence. **(B)** The same analysis is presented for mutations identified from individually collected mutant plaques. Whereas Duplex Sequencing recovers all functional classes of predicted amino acid mutations along the entire gene, mutations from picked mutant plaques that have lost a functional cII protein are devoid of synonymous variants and mutations at the non-essential C- and N-termini. Nucleotide positions with higher than average mutation counts by Duplex Sequencing reflect mutagenic hotspots. The different mutation profile observed in the TGR plaque sequencing is more reflective of which sites are most phenotypically selected.

### Rates of chemical-induced mutagenesis vary by genomic locus

TGR assays rely on the assumption that the mutability of the *cII* lambda phage transgene is a representative surrogate for the entire mammalian genome. We hypothesized that local genomic features and functions of the genome such as transcriptional status, chromatin structure, and sequence context may modulate mutagenic sensitivity.

To test this idea, we used DS to measure the exposure-induced spectrum of mutations in 4 endogenous genes with different transcriptional status in different tissues: beta catenin *(Ctnnb1),* DNA-directed RNA polymerases I and III subunit RPAC1 *(Polr1c),* haptoglobin *(Hp)* and rhodopsin (*Rho*), as well as the *cII* transgene in Big Blue mouse liver and marrow of animals exposed to olive oil (vehicle control), B[α]P, or ENU. We assessed mutations in the same four endogenous loci in the lung, spleen, and blood of Tg-rasH2 mice exposed to saline (vehicle control) or urethane to investigate DS performance in a second mouse model.

The Duplex Sequencing single nucleotide variant (SNV) per-nucleotide mutant frequencies across mouse model, tissue, treatment group, and genomic locus are shown in **Figure 4**. Vehicle control (VC) mutant frequencies averaged 1.14× 10^−7^ in the Big Blue mouse model (**Fig. 4A**) and 9.03 × 10^−8^ in the Tg-rasH2 mouse model (**Fig. 4b**). The number of unique mutant nucleotides detected per VC sample ranged from 5 to 36 (mean 15.5) and were always non-zero (**Fig. S3**.) These frequencies are comparable to Chawanthayatham *et al.* (2017) where a DMSO vehicle-exposed transgenic *gptΔ* mouse was measured to have a mutant frequency of 2.7×10^−7^ in liver samples after Duplex Sequencing of the reporter recovered from gDNA^28^. We observed a mean background mutant frequency in the marrow (1.63 × 10^−7^) nearly twice that of peripheral blood (1.06× 10^−7^), liver (9.63 ×10^−8^), lung (7.13 ×10^−8^), and spleen (7.45 ×10^−8^), which may relate to differences in relative cell-cycling times in these tissues.

**Figure 4.**
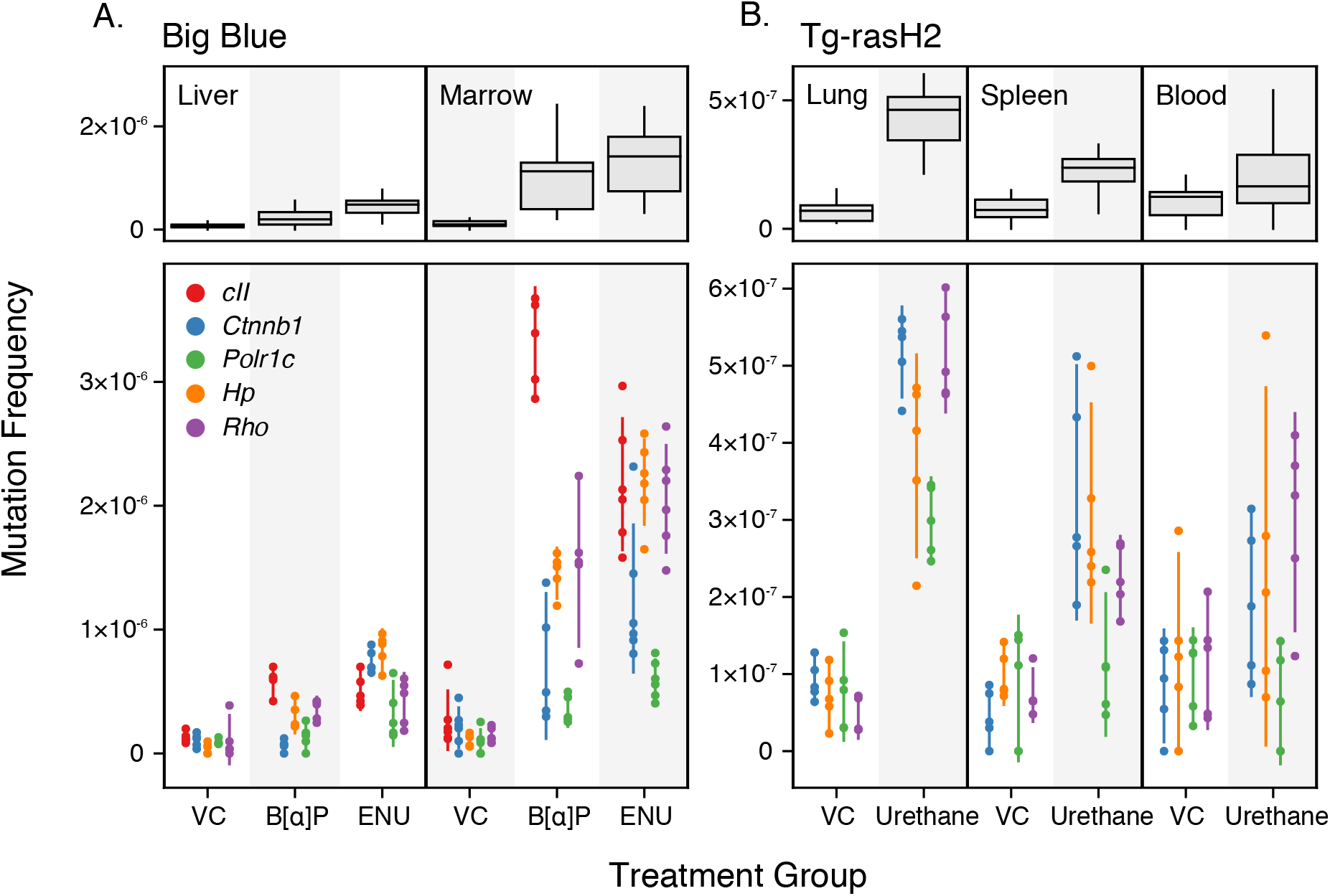
Sensitivity to mutagenesis varies by tissue type, mutagen, and genomic locus. Single nucleotide variant (SNV) per-nucleotide mutant frequency (MF) is shown by tissue and treatment aggregated across all loci interrogated (top) and by individual genic regions (bottom). Box plots show all four quartiles of all data points for that tissue and treatment group. Scatter points show individual MF measurements from replicate animals in each cohort with line segments representing 95% CI. **(A)** Big Blue mouse study evaluating liver and bone marrow in animals exposed to vehicle control (VC), B[α]P, or ENU. **(B)** Tg-rasH2 mouse study evaluating lung, spleen, and peripheral blood in animals exposed to VC or urethane. There is no *cII* transgene in the Tg-rasH2 mouse model. Note the different Y-axis scaling between the two studies.

In all mutagen-exposed samples, the mutant frequency was increased over the respective vehicle control samples. However, the fold-induction across tissue types varied considerably, as each compound has a different mutagenic potential, presumably related to varying physiologic factors such as tissue distribution, metabolism, and sensitivity to cell-turnover rate^29,30^.

The *cII* and *Rho* genes had highest mutant frequencies among all tested loci in bone marrow. Other genes, such as *Ctnnb1* and *Polr1c,* exhibited frequencies as much as 8-fold lower. This disparity is potentially due to the differential impact of transcription levels and transcription-coupled repair (TCR) of lesions and/or local chromatin structure^31^. *Ctnnb1* and *Polr1c* are thought to be transcribed in all tissues we tested, and therefore benefit from TCR, whereas *Rho* and *cII* are thought to be non-transcribed, and thus should not be impacted by TCR.

*Hp* was selected as a test gene because it is transcribed in the liver but not significantly in other tissues. The aforementioned logic cannot explain why *Hp* exhibited an elevated mutation rate compared to other genomic loci in the mouse liver. An additional genomic process related to the transcriptional status is DNA methylation. It is known that lesions on nucleotides immediately adjacent to a methylated cytosine have a lower probability of being repaired due to the relative bulk and proximal clustering of the adducts^32^. This or other factors, such as differential base composition between sites could also be at play.

Mechanisms aside, the widely variable mutant frequency we observe across different genomic loci indicates that no single locus is ever likely to be a comprehensive surrogate of the genome-wide impact of chemical mutation induction.

### Strand bias of mutations reflect functional effects of the genome

To further investigate the potential role of TCR as a contributor to the observed differential regional sensitivity to mutagens, we examined the strandedness of mutations identified by Duplex Sequencing at each locus. Mutational strand bias is defined as a difference in the relative propensity for a particular type of nucleotide change *(e.g.* A→C) to occur on one DNA strand versus the other. This bias may result from multiple factors including transcription, epigenetic influences (*e.g.* methylation), proximity to replication origins, and nucleotide composition, among others^33,34^. We compared the per-nucleotide mutant frequency for each base substitution against its reciprocal substitution in our urethane-exposed mouse cohort. If a strand bias were to exist, then these frequencies would be unequal^35^. We then correlated the extent of strand differences observed by genic region with predicted transcriptional status of each tissue.

Human transcription levels of four genes *(Ctnnb1, Polr1c, Hp, Rho)* were used as a surrogate for those in mouse tissues and were obtained from the Genotype-Tissue Expression (GTEx) Project Portal (accessed on 2020-01-06). In humans, the levels of *Ctnnb1* expression are highest in lung (median transcripts per million [TPM] 164.4) and lower in spleen and blood (median TPM 100.3 and 25.75 respectively) whereas levels of *Polr1c* expression are low in all three (median TPM 19.27,24.09 and 3.83, respectively). In humans, the genes *Hp* and *Rho* are largely nonexpressed in spleen, lung, and blood.

Two genomic regions, *Ctnnb1* and *Polr1c,* showed high urethane-mediated strand bias (**Fig. 5**), which is consistent with a model of transcription-coupled repair since TCR predominantly repairs lesions on the transcribed strands of active genes^33^. The majority of observed strand bias fell into two base substitution groups (T·A→A·T and T·A→G·C) in genes expressed in lung tissue *(Ctnnb1* and *Polr1c).* The mean reciprocal SNV fold-difference of these mutation types across all tissue types was 11.6 and 9.0 in *Ctnnb1* and *Polr1c* versus 1.6 and 0.8 in (non-transcribed) *Hp* and *Rho*. The highest bias existed in lung tissue which is consistent with a TCR-related mechanism given that lung has the highest predicted transcription rate among the tissue types assayed.

**Figure 5.**
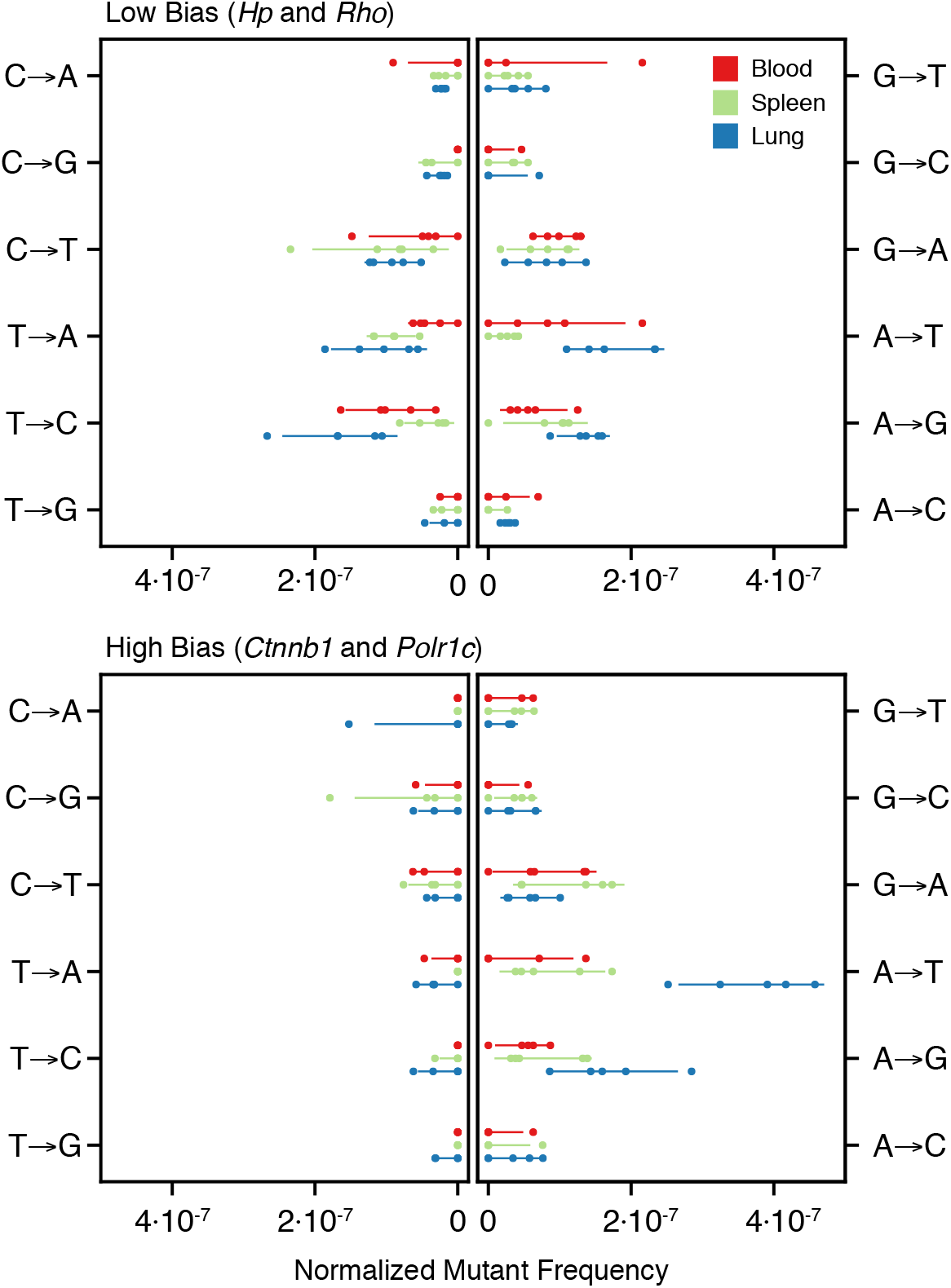
Strand bias in base substitutions exists in regions of moderate to high transcription. Mutational strand bias was seen for urethane exposed tissues in *Ctnnb1* and *Polr1c* (expressed in tissues examined) but not in *Hp* or *Rho* (not expressed in tissues examined). Single nucleotide variants (SNV) are normalized to the reference nucleotide in the forward direction of the transcribed strand. Individual replicates are shown with points, and 95% confidence intervals with line segments. Mutant frequencies were corrected for the nucleotide counts of each reference base in the target genes. The observed bias is evident in *Ctnnb1* and *Polr1c* as elevated frequencies of A→N and G→N variants relative to their complementary mutation *(i.e.* asymmetry around the vertical line), in contrast to the balanced spectrum of *Hp* and *Rho.* This difference is likely due to the mutationattenuating effect of transcription-coupled repair on the template strand of transcribed regions of the genome.

### Unsupervised clustering resolves simple patterns of mutagenesis

We next sought to classify each sample into a mutagen class based solely on the simple spectrum of single nucleotide variants observed within the endogenous regions examined in both the Big Blue and Tg-rasH2 animals. The technique of unsupervised hierarchical clustering can resolve patterns of spectra as distinct clusters with common features^36^. **Figure 6A** shows a strong spectral distinction between ENU and both VC and B[α]P. However, the simple spectra of VC and B[α]P resolve poorly. A gradient of similarity is apparent in the VC and B[α]P cluster which suggests that, with deeper sequencing, it may be possible to fully resolve the two. No statistically valid clusters emerged that correlates with tissue type, suggesting that the patterns of mutagenesis for both B[α]P and ENU are similar in the liver and marrow of the Big Blue mouse. **Figure 6B** shows perfect clustering by exposure due to the orthogonal patterns of urethane mutagenesis as compared to the unexposed tissues in Tg-rasH2 mice. We similarly saw no correlated clustering at the level of tissue type in Tg-rasH2 mice.

**Figure 6.**
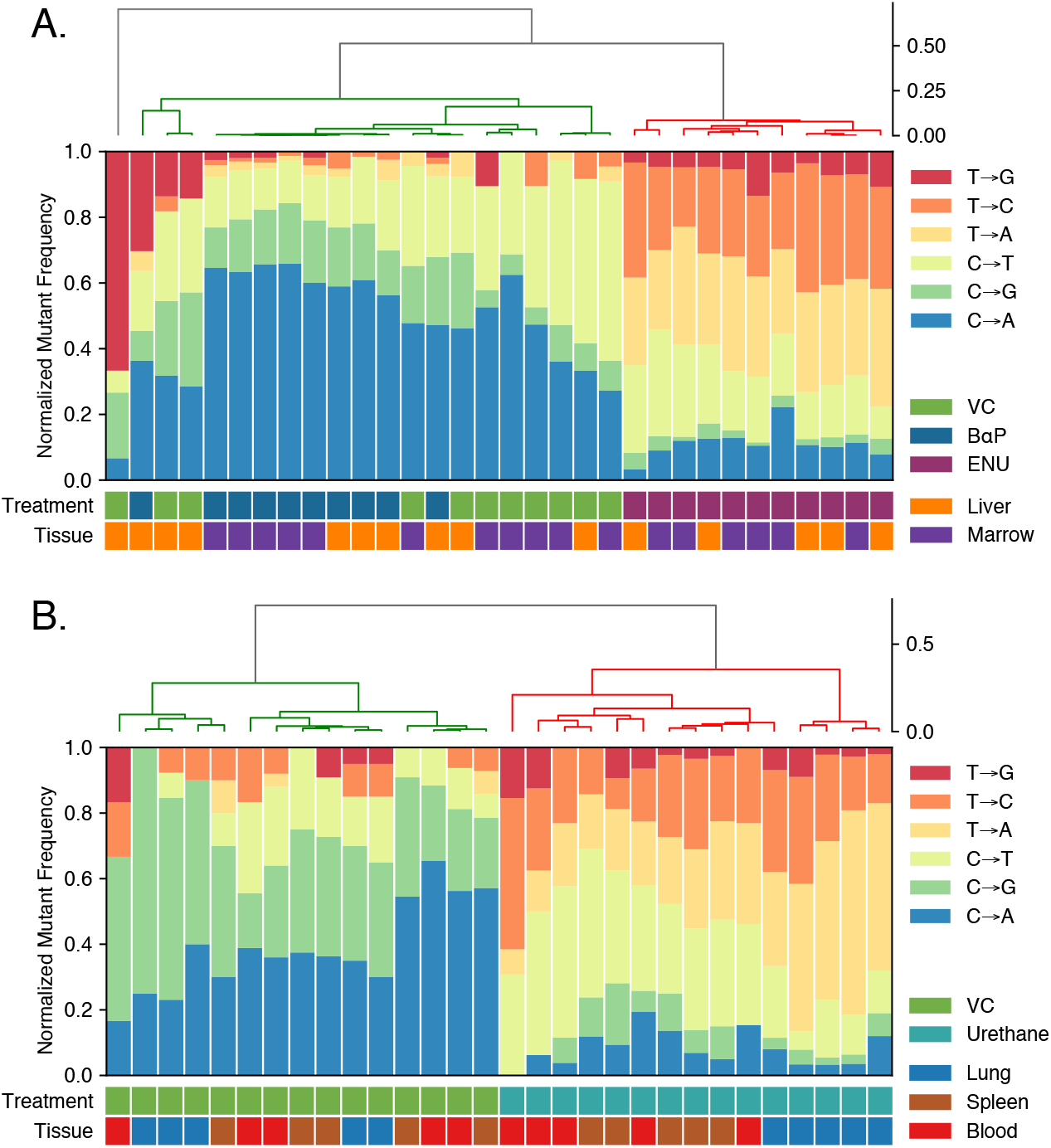
Unsupervised hierarchical clustering predicts mutagen treatment across samples. Clustering of simple spectrum probabilities was performed with the weighted (WGMA) method and cosine similarity metric. **(A)** Liver and marrow in Big Blue animals exposed to ENU, B[α]P, or vehicle control (VC). **(B)** Lung, spleen, and blood samples from the Tg-rasH2 cohort exposed to urethane or VC. Clustering was near-perfect except for distinguishing B[α]P from vehicle exposure in liver tissue where fewer mutational events were observed due to its lower proliferation rate.

### Trinucleotide spectrum of treatment groups show distinct patterns of mutagenesis and relate to patterns seen in human cancer

To further classify the patterns of single nucleotide variants by treatment group, we considered all possibilities of the 5’ and 3’ bases adjacent to the mutated base to create trinucleotide spectra^12,36,37^. When enumerating all 96 possible single nucleotide variants within a unique trinucleotide context, a distinct pattern for each treatment group becomes apparent (**Fig. 7 A-D**) that show similarities to mutational signatures as extracted from thousands of human cancers (**Fig. 7E**).

**Figure 7.**
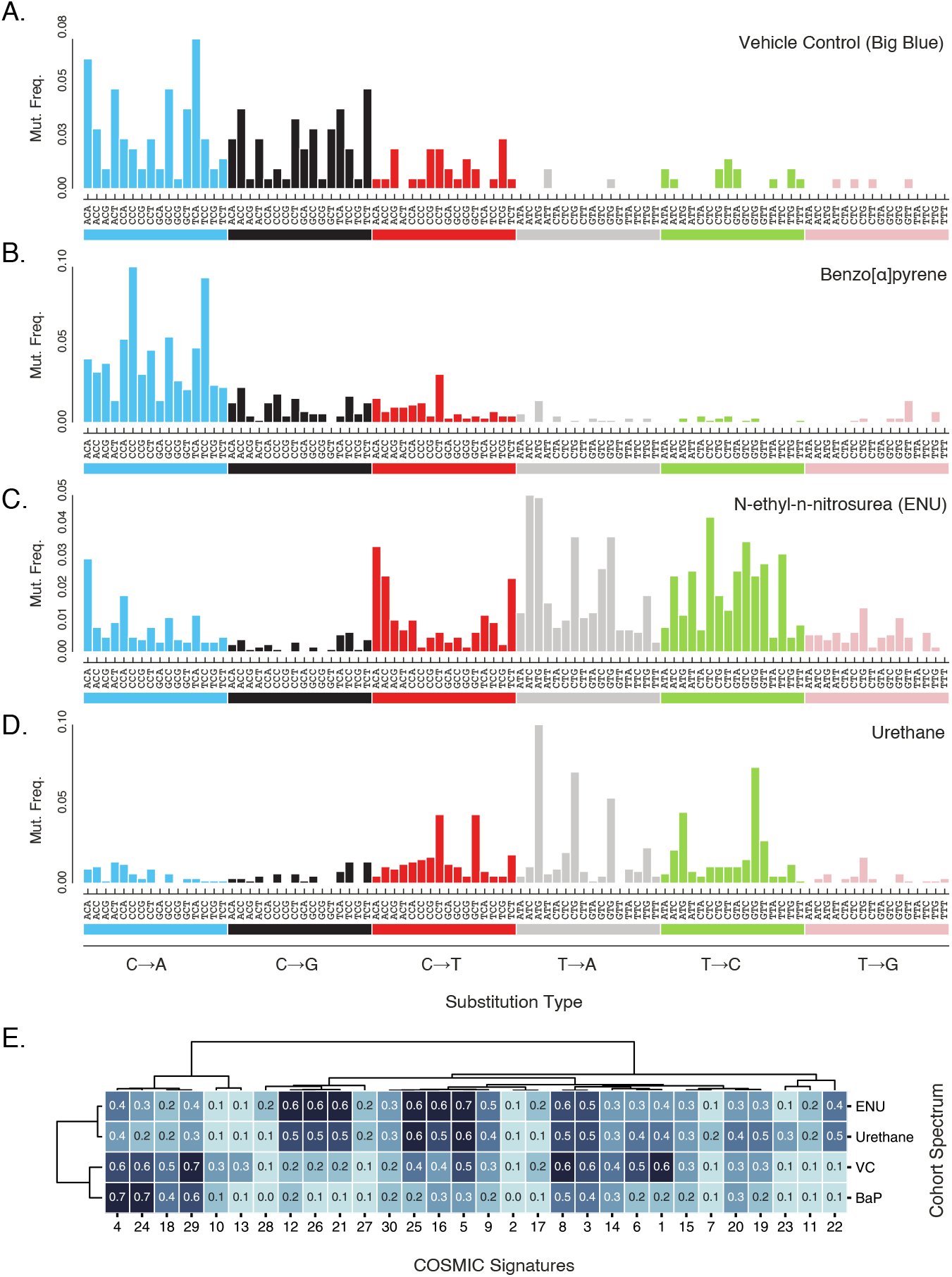
Trinucleotide base substitution spectra of each mutagen treatment reflects distinct mutational processes. The proportion of base substitutions in all trinucleotide contexts (pyrimidine notation) for the union of all endogenous mouse genic regions in **(A)** vehicle control **(B)** benzo[α]pyrene, **(C)** N-ethyl-N-nitrosourea and **(D)** exposed mice. Each proportion was derived by normalizing the observed substitution types in each context by the relative abundance of that context in the regions examined. **(E)** Unsupervised hierarchical clustering of the first 30 published COSMIC signatures and the 4 cohort spectra. Clustering was performed with the weighted (WGMA) method and cosine similarity metric. B[α]P is most similar to Signatures 4, 24, and 29. Signature 24 is correlated with Aflatoxin B1 exposure and has a similar mutagenic mode of action to the DNA intercalating reactive metabolites of B[α]P. Vehicle control (VC) is most like Signature 1, which is believed to reflect the age-associated mutagenic effect of reactive oxidative species and deamination.

The VC trinucleotide spectrum (**Fig. 7A**) is most similar to Signature 1 from the COSMIC catalog of somatic mutation signatures in human cancer^38^ (cosine similarity of 0.6), which has a proposed etiology of C·G→T·A transitions in CpG sites resulting from unrepaired spontaneous deamination events at 5-methyl-cytosines. The most notable difference between the bulk trinucleotide spectrum of VC and Signature 1 is the extent of C·G→A·T and C·G→G·C transversions which most likely reflect endogenous oxidative damage, an age-related process^39^.

The B[α]P trinucleotide spectrum (**Fig. 7B**) is predominantly driven by C·G→A·T mutations with a higher affinity for CpG sites. This observation is consistent with previous literature indicating that B[α]P adducts, when not repaired by TCR, lead to mutations most commonly found in sites of methylated CpG dinucleotides^32,37^ This spectrum is highly similar to Signature 4 (0.7 cosine similarity) and Signature 29 (0.6 cosine similarity), both of which have proposed etiologies related to human exposure to tobacco where B[α]P and other polycyclic aromatic hydrocarbons are major mutagenic carcinogens. The spectrum for *in vivo* murine exposure to B[α]P is equally comparable to Signature 4 and Signature 24 (0.7 cosine similarity), likely due to similar mutagenic modes of action between B[α]P and aflatoxin (the proposed etiology of signature 24)^36^.

The urethane trinucleotide spectrum (**Fig. 7D**) has no confidently assignable analog in the COSMIC signature set. As compared to the simple spectrum of urethane in **Figure 6B**, a periodic pattern of T·A→A·T in 5’-NTG-3’ emerges. This pattern of highly residue-specific mutagenicity has been previously observed in the trinucleotide spectra of whole genome sequencing data from adenomas of urethane-exposed mice^40^ as well as, very recently, in urethane-exposed lung tissue of mice weeks after exposure, as detected by another ecNGS method^41^.

### Oncogenic Ras mutations undergo strong *in vivo* selection within weeks of carcinogen exposure in cancer-prone Tg-rasH2 mice

The Tg-rasH2 mouse model contains 4 tandem copies of human *HRAS* with an activating enhancer mutation to boost oncogene expression^13^. The combination of enhanced transcription and boosted proto-oncogene copy number predisposes the strain to cancer. Use of these mice in a 6-month cancer bioassay is accepted under ICH S1B guidelines as an accelerated substitute for the traditional 2-year mouse cancer bioassay used for pharmaceutical safety assessment^42^. Exposure to urethane, a commonly used positive control mutagen, results in splenic hemangiosarcomas and lung adenocarcinomas in nearly all animals by 10 weeks post-exposure.

We examined the effect of urethane exposure on the *HRAS* transgene, as well as the endogenous *Hras, Kras,* and *Nras* genes, at DNA residues most commonly mutated in human cancers (**Fig. 8**).

**Figure 8.**
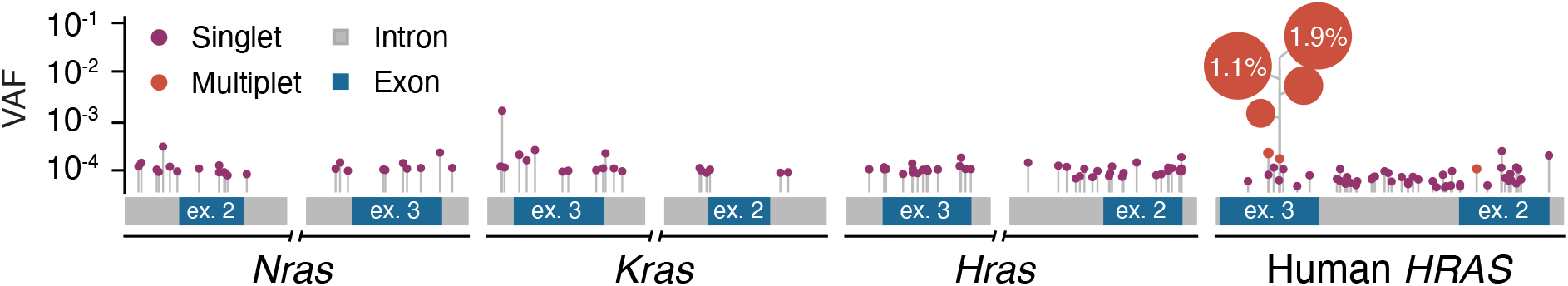
Early neoplastic evolution in cancer-prone mice following carcinogen exposure. The location and variant allele frequency (VAF) of single nucleotide variants are plotted across the genomic intervals for the introns and exons captured from the endogenous mouse *Ras* family of genes as well as the human transgenic *HRAS* loci from the Tg-rasH2 mouse model. Singlets are mutations identified in a single molecule of a sample. Multiplets are an identical mutation identified within multiple molecules within the same sample and may represent a clonal expansion event. Pooled data from all tissues in the experiment (lung, spleen, blood) are included. The height of each point (log scale) corresponds to the VAF of each single nucleotide variant (SNV). The size of the point corresponds to the number of counts observed for the mutant allele. A cluster of multiplet A·T →T·A transversions at the human oncogenic *HRAS* codon 61 hotspot is seen in 4 out of 5 urethane exposed lung samples and 1 out of 5 urethane exposed splenic samples (**Supplemental Table 4**). The observation of an identical mutation in independent samples with high frequency multiplets in a well-established cancer driver gene likely indicates positive selection. Notably, these clones are defined by the transversion A·T →T ·A in the context NTG which is characteristic of urethane mutagenesis.

In contrast to the endogenous *Ras* family genes, the human *HRAS* transgene is present in four copies per haploid genome–each under the control of a tandem promoter/enhancer, but without the repression system that is present at the native human *HRAS* locus. We postulated that the mechanism of activation of human *HRAS* in the Tg-rasH2 model would positively influence selection of the cells harboring the activating mutations and would be observable as outgrowth of clones bearing mutations at hotspot residues relative to residues not under positive selection. Indeed, we observed compelling signs of selection, as evidenced by focally high variant allele frequencies (VAF) of activating mutations at the canonical codon 61 hotspot in exon 3 in the human *HRAS* transgene, but not at other sites in that gene, nor at homologous sites in the endogenous mouse *Ras* family. Sizable clonal expansions of this mutation were detected in 4 out of 5 lung samples, 1 out of 5 spleen samples, and in no blood samples which is consistent with the historically known relative frequency of tumors in each tissue.

Moreover, not only are the variant allele frequencies as much as 100-fold higher than seen for any other endogenous gene variant, but the absolute counts of mutant alleles at this locus is very high (>5), which offers strong statistical support for these clones existing as authentic expansions and not as independent mutated residues occurring by chance (**Supplemental Table 4**). Notably, all clonal mutations observed at codon 61 are A·T→T·A transversions in the context 5’-CTG-3’, which conforms to the context 5’-NTG-3’ (where N is the IUPAC code for any valid base), which is highly mutated across all genes in the urethane exposed mouse samples^40^ (**Fig 6D**). Other types of mutations at codon 61 could lead to the same amino acid change, so the combination of the specific nucleotide substitution observed, the clone size relative to that of other loci, and the repeated observation across independent samples of the most tumor prone tissues paints a comprehensive picture of both a urethane-mediated mutagenic trigger and a carcinogenic process that follows.

## DISCUSSION

We have demonstrated that Duplex Sequencing, an extremely accurate error-corrected NGS (ecNGS) method, is a powerful tool for the field of genetic toxicology that can be used to assess both mutagenesis and carcinogenesis *in vivo*. Unlike conventional *in vivo* mutagenesis assays, Duplex Sequencing does not rely on selection, but rather on unbiased digital counting of billions of individual nucleotides directly from the DNA region of interest. This yields data that is both richer and more broadly representative of the genome than current tools and allows fundamentally new types of biological questions to be asked.

From sequence data it is possible to mine a wealth of information including mutation spectrum, trinucleotide mutation signatures, and predicted functional consequences of mutations. By virtue of not being limited to a specific reporter, we showed that the relative susceptibility to chemical mutagenesis varies significantly by genomic locus and is further influenced by tissue. We could infer this to be (at least partially) the result of non-uniform transcription-coupled repair, as evidenced by the consistent asymmetry of certain mutation types between transcribed and nontranscribed strands. The examples shown here are limited by the modest number of loci and tissues and the inference of transcriptional status based on another species and can be improved upon in future studies. It is likely that many other factors beyond transcriptional status shape the relative plasticity of the genome and can be uncovered with careful investigations.

The ability to directly observe subtle regional mutant frequency differences, on the order of one-in-ten million, is extraordinary in terms of biological study opportunities, but also raises practical questions for regulatory usage. For example, what would define the optimal subset of the genome to be used for drug and chemical safety testing? For some applications, a diverse, genome-representative panel makes the most sense; for others it might be preferable to enrich for regions that are predisposed to certain mutagenic processes^43^ or have unique repair biology^35^.

Not all carcinogens are mutagens. Those which are not mutagenic will not produce a signal in mutagenesis assays–either conventional or sequencing-based. However, as shown here, it appears possible to use ecNGS to infer carcinogenesis via detection of clonal expansions carrying oncogenic driver mutations as a marker of a neoplastic phenotype^44^ This concept is more complex to design, insofar as it necessitates some *a priori* knowledge about the common drivers that are operative in different tissues in response to different classes of carcinogens. However, there is simply no other approach, convenient or not, that can quantitate these signals in less than a month from exposure. The proof-of-concept illustrated here relied on a mutagenic chemical in a cancer-predisposed mouse strain; future efforts will be needed to demonstrate the same with non-genotoxic carcinogens in wild-type animals.

A further advantage of ecNGS is the breadth of applicability, *in vivo* or *in vitro,* to any tissue from any species. *In vivo* selection-based assays are organism and reporter specific; the former restricts testing to rodents, and the latter confers potential biases to mutational spectrum and does not allow targeting of specific genomic regions. The only *in vivo* mutagenesis assay that does not depend on *in vitro* selection, the Pig-a Gene Mutation Assay, classically restricted to only erythrocytes, requires bioavailability to the bone marrow compartment, cannot be used for spectrum analysis, and necessitates access to flow cytometery equipment^45^. In contrast, next generation DNA sequencing platforms are widely available and can be automated to handle thousands of samples per day, thus rendering the approach tractable for many different types of labs.

We are not the first to apply NGS to mutagenesis applications^12^. Sequencing the reporter gene from pooled clones from TGRs has been used to identify *in vivo* mutagenic signatures^17^. Single cell cloning of mutagen-exposed cultured cells and patient-derived organoids has been used to identify *in vitro* and *in vivo* mutagenic signatures^46,47,48^. In each case, cloning, followed by biological amplification was required to resolve single-cell mutational signals, which would otherwise be undetectable in a background of sequencing errors. We have previously used Duplex Sequencing to measure trinucleotide signatures in phage-recovered reporter DNA of mutagen-exposed transgenic mice without the need for cloning^36^. Others have characterized mutational spectra directly from human DNA using a form of very low depth, whole genome, Duplex Sequencing without added molecular tags^49^. However, each of these methods has factors that limit its practicality for broad usability.

The cost of any NGS-based technique is an important consideration, particularly when compared to something as routine as the bacterial Ames assay. Duplex Sequencing further multiplies sequencing costs because of the need for multiple redundant copies of each source strand as part of the consensus-based error correction strategy. However, over the last 12 years, the cost of NGS has fallen nearly 4 orders of magnitude, whereas the cost of conventional genetic toxicology assays has remained largely unchanged. Extrapolating forward, we anticipate that equipoise will eventually be reached. Savings by virtue of not needing to breed genetically engineered animals, the ability to repurpose tissue or cells already generated for other assays (supporting the 3R concept of replacement, reduction, and refinement^50^), decreased labor, and greater automatability should also serve to increase efficiencies and lessen animal use.

Beyond being undesirable, animal testing is simply not possible for some applications. New forms of mutagenesis, such as CRISPR and other gene editing technologies, are highly sequence-specific and cannot be easily de-risked in alternative genomes or using reporter genes^51,52^. Being able to carry out rapid in-human genotoxicity assessment as a part of early clinical trials may also be important for applications where there is urgency to develop therapies, such as drugs being tested against the 2019 pandemic coronavirus^53^ and those needed in future public health emergencies.

Controlled drug and chemical safety testing are not the only reasons to screen for mutagenic and carcinogenic processes. Humans are inadvertently exposed to many environmental carcinogens^54,55^. The ability to identify biomarkers of mutagenic exposures using DNA from tissue, or non-invasive samples such as blood, urine, or saliva is an opportunity for managing individual patients via risk-stratified cancer screening efforts as well as public health surveillance to facilitate carcinogenic source control^12^. Deeper investigations into human cancer clusters^56^, monitoring those at risk of occupational carcinogenic exposures^57^, such as firefighters^58^ and astronauts^59^, and surveilling the genomes of sentinel species in the environment as first-alarm biosensors^60^ are all made possible when DNA can be analyzed directly.

Almost four decades have passed since it was envisioned that the entirety of one’s exposure history might be gleaned from a single drop of blood^61^. While this remains a lofty ambition, the data we have shown here suggest that it is not wholly implausible. Our work indicates that there is a much greater amount of information recorded in the somatic genome than we have previously been able to appreciate, or access. Future studies are needed to determine how best to capitalize on this data for basic research applications, preclinical safety testing, and in-human studies.

## MATERIALS AND METHODS

### Animal Treatment and Tissue Collection

All animals used in this study were housed at AAALAC international accredited facilities and all research protocols were approved by these facilities respective to their Institutional Animal Care and Use Committees (IACUC).

Big Blue^®^ C57BL/6 homozygous male mice [C57BL/6-Tg(TacLIZa)A1Jsh] bred by Taconic Biosciences on behalf of BioReliance were dosed daily by oral gavage with 5 mL/kg vehicle control (olive oil) or benzo[α]pyrene (B[α]P) formulated in the vehicle at a dose level of 50 mg/kg per day for 28 days. A third cohort of Big Blue mice were dosed by oral gavage with 40 mg/kg per day (10 mL/kg) of N-ethyl-N-nitrosourea (ENU) formulated in phosphate buffer solution (pH 6.0) on days 1, 2 and 3. All animals were necropsied on study day 31.

Tg-rasH2 male mice [CByB6F1-Tg(HRAS)2Jic] from Taconic Biosciences received a total of three intraperitoneal injections of vehicle control (saline) or urethane (1000 mg/kg per injection) at a dose volume of 10 mL/kg per injection on days 1, 3 and 5. Animals were necropsied on study day 29.

Liver, lung, and spleen samples were collected and then flash frozen. Bone marrow was flushed from femurs with saline, centrifuged, and the resulting pellet was flash frozen. Blood was collected in K2 EDTA tubes and flash frozen.

Studies were generally consistent with OECD TG 488 guidelines except that ENU and urethane were dosed less than daily but at a frequency known to produce systemic mutagenic exposures. The sampling time for the urethane study was at day 29 and not day 31.

### Plaque Assay for Mutant Analysis

High molecular weight DNA was isolated from frozen Big Blue and Tg-rasH2 tissues using methods as described in the RecoverEase product use manual Rev. B (Cat # 720202, Agilent Inc, Santa Clara CA, USA). Vector recovery from genomic DNA, vector packaging into infectious lambda phage particles, and plating for mutant analysis was performed using methods described in the λ Select-cII Mutation Detection System for Big Blue Rodents product use manual Rev. A (Cat # 720120, Agilent, Santa Clara CA, USA)^5^.

### Phage and Mouse DNA for Duplex Sequencing

Phage DNA was purified from phage plaques punched from the *E. coli* lawn on agar mutant selection plates following 2 days of incubation at 24° C. Agar plugs were pooled by mutagen treatment group in SM buffer and then frozen for storage. DNA was purified using the QIAEX II Gel Extraction Kit (Cat # 20021, Qiagen, Hilden, Germany). Mouse genomic DNA was purified from liver, bone marrow, lung, spleen, and blood. Approximately 3 mm x 3 mm x 3 mm tissue sections were pulverized with a tube pestle in a microfuge tube. DNA was extracted using the Qiagen DNeasy Blood and Tissue Kit (Cat # 69504, Qiagen, Hilden, Germany).

### Duplex Sequencing

Extracted genomic DNA was ultrasonically sheared to a median fragment size of approximately 300 base pairs using a Covaris system. Sheared DNA was further processed using a prototype cocktail of enzymes with glycosylase and lyase activity for the purpose of excising certain forms of DNA damage and cleaving phosphodiester backbones at resulting abasic sites to render damaged, or incomplete duplex templates un-amplifiable (TwinStrand Biosciences, Seattle WA, USA). DNA was end-polished, A-tailed, and ligated to Duplex Sequencing Adapters containing semi-degenerate unique molecular identifies (TwinStrand Biosciences, Seattle WA, USA) via the general method described previously^9,16^. Adapter-ligated DNA fragments were then PCR amplified with primers containing dual unique indexes. After the initial PCR, samples were individually subjected to tandem hybrid capture using 120mer biotinylated DNA oligo probes (IDT, Coralville, IA, USA), for a total of two captures. The first (indexing) and second PCR respectively entailed 10 and 14 cycles. The third PCR involved a variable number of cycles until the library could be accurately quantified. Resulting libraries were quantified, pooled, and sequenced on an Illumina NextSeq 500 using 150 base pair paired-end reads with vendor supplied reagents. Where necessary, SYBR-based qPCR was used to determine appropriate DNA input by normalizing phage and mouse DNA across library preparations by total genome equivalents. Library input, before shearing, of plaque DNA was ~100 pg and the genomic DNA input for all mouse samples was ~500 ng. A summary of sequencing data yields for Big Blue and Tg-rasH2 samples is listed in **Supplemental Tables 2 and 3**.

### Hybrid Selection Panel Design

Hybrid selection baits for all targets were designed to intentionally avoid capturing any nucleotide sequence within 10 bp of a repeat-masked interval as defined in RepBase (**Fig. S4**)^62^. Intronic regions adjacent to the exons of the target genes were baited to provide a functionally neutral and non-coding view on the pressures of mutagenesis near exonic targets. Duplex consensus base pairs and subsequent variant calls were only reported over a region defined by the same repeat-mask rule as for the bait target design. All libraries achieved 99.9% alignment of duplex consensus bases over the target territories with less than 0.001% of off-panel alignment. All targets were of expected uniform coverage given that no off-target alignment to pseudo-genes or repetitive genomic sequences was observed.

Baits were also designed to target the *cII* transgene in the Big Blue mouse model and the human *HRAS* transgene in the Tg-rasH2 mouse model. The multi-copy *cII* transgene was sequenced to a median target coverage of 39,668x and the multi-copy human *HRAS* transgene to 9,012x.

### Consensus calling and consensus post-processing

Consensus calling was carried out as described in “Calling Duplex Consensus Reads” from the Fulcrum Genomics fgbio tool suite^63^. Generally, the algorithm proceeds with aligning the raw reads with bwa. After alignment, read pairs were grouped based on the corrected unique molecular identifier (UMI) nucleotide bases and their shear point pair as determined through primary mapping coordinates. The read pairs within their read pair groups were then unmapped and oriented into the direction they were in as outputted from the sequencing instrument. Quality trimming using a running-sum algorithm was used to eliminate poor quality three-prime sequence. Bases with low quality were masked to ‘N’ for an ambiguous base assignment. Cigar filtering and cigar grouping was performed within each read pair group to help mitigate the poisoning effect of artifactual indels in individual reads introduced in library preparation or sequencing. Finally, consensus reads were created, from which duplex consensus reads meeting pre-specified confidence criteria were filtered. Barcode error-correction was performed using a known whitelist of barcodes, a maximum number of mismatches between a barcode and an expected barcode of 1, and a minimum Hamming distance to the next most likely known barcode of 2. After duplex consensus calling, the read pairs underwent balanced overlap hard clipping to eliminate biases from double counting bases due to duplicate observation within an overlapping paired-end read. Duplex consensus reads were then end-trimmed and inter-species decontamination was performed using a *k*-mer based taxonomic classifier (**Supplementary Methods**).

### Variant calling and variant interpretation

Variants were called using VarDictJava with all parameters optimized to collect variants of any alternate allelic count greater than, or equal to, one^64^.

There are two polar interpretations one can make when an identical canonical variant is observed multiple times in the same sample. The first assumption is that the observations were independent; that they were acquired during unrelated episodes in multiple independent cells and are not the product of a clonal expansion and shared cell lineage. The second assumption is that the alternate allele observations are a clonal expansion of a single mutagenic event and can all be attributed to one initial mutagenic event.

When classifying variant calls as either independent observations, or from a clonal origin, we first fit a long-tailed distribution to all variants that were not germline. Any outliers to this distribution with multiple observations are deemed to have arisen from a single origin. This method may serve to undercount multiple independent mutations at the same site under extraordinary specific mutagenic conditions. For example, the clonally expanded A·T→T·A transversion at codon 61 in the *HRAS* transgene was a significant outlier to this model and was highly correlated with urethane exposure. The variant allele frequency (VAF) of these expanded mutations varied 100x in urethane-exposed lung tissues, however, our calculation of per-nucleotide mutant frequency varied only ~2x indicating that this one tissue-specific residue was under the highest selective pressures for expansion beyond any other residue in any other tissue within the panel territory.

### Hierarchical clustering of base substitution spectra

All clustering was performed using the Wald method and the cosine distance metric. Leaves were ordered based on a fast-optimal ordering algorithm^65^. Simple base substitution spectrum clustering was achieved by first normalizing all base substitutions into pyrimidine-space and then normalizing by the frequencies of nucleotides in the target region. Clustering of trinucleotide spectra was achieved in a similar manner where base substitutions were normalized into pyrimidine-space and then partitioned into 16 categories based on all the combinations of five and three prime adjacent bases^38^. Subsequent normalization of trinucleotide spectra was performed using the frequencies of 3-mers in the target regions.

## Supporting information

Database S1

Database S2

## ABBREVIATIONS

B[α]P: Benzo[α]pyrene
DS: Duplex Sequencing
ecNGS: Error-Corrected Next Generation Sequencing
ENU: N-Ethyl-N-Nitrosourea
MF: Mutant Frequency
SNV: Single Nucleotide Variant
TCR: Transcription-Coupled Repair
TGR: Transgenic Rodent
TPM: Transcripts per million
VC: Vehicle Control
VAF: Variant Allele Frequency

## Acknowledgements

This work was partially funded by NIH R44 ES030642 to JJS. Tg-rasH2 mice were kindly provided by Taconic Biosciences, Inc., Germantown, NY, US. Marie McKeon of MilliporeSigma was responsible for the experiments during the in-life phase of the Tg-rasH2 study. The Genotype-Tissue Expression (GTEx) Project database used for transcript level approximation was supported by the Common Fund of the Office of the Director of the National Institutes of Health, and by NCI, NHGRI, NHLBI, NIDA, NIMH, and NINDS. The authors wish to thank others at TwinStrand Biosciences, Amgen, MilliporeSigma, members of the HESI GTTC consortium and Dr. Larry Loeb for intellectual support throughout these studies.

## Data and materials availability

Final filtered and decontaminated error-corrected alignments for all 62 mouse samples in the BAM file format are deposited in the Sequence Read Archive under BioProject accession PRJNA673916.

## Author Contributions

CCV and JJS wrote the manuscript. CCV analyzed the data. CCV, RRY, MRF, SM, RK, LNW, TL, and JJS designed and performed the experiments and provided review of the manuscript.

## SUPPLEMENTARY METHODS

### Inter-species contamination is robustly detected by Duplex Sequencing

A consequence of extremely accurate error-correction next generation sequencing (ecNGS) technologies is the detection of ultra-rare intra-species contamination and how false positive alignments of those sequencing reads can bias per-nucleotide mutant frequency (MF) calculation by more than 100-fold. The false positive alignment of short reads not from the target species is particularly likely when sample processing is done near samples that are of an alternate species. This issue is exacerbated when targeting regions of high homology among all species, such as in conserved or exonic regions of the genome (**Fig. S5**).

The solution we developed for handling inter-species read-pair decontamination relies on taxonomic classification of all error-corrected sequences from the entire study to ensure only the read pairs that match the target species with high confidence are kept for downstream analysis.

A taxonomy database was constructed with *k*-mers from human, rat, cow, and mouse. The taxonomic classifier Kraken^66^ was used to identify error-corrected paired-end contaminating reads, as well as confidently indicating which reads were only from *Mus musculus* origin. Reads that are left unassigned due to this method are often true sequences from the source genomes, however, they contain an ‘N’-call or variant base often enough such that a single *k*-mer cannot exist that indicates a positive classification to the target genome. Reads of ambiguous assignment were discarded as they did not contain enough sequence information to positively assign them to any of the organisms at the species level.

To eliminate confounding assignment due to the human *HRAS* transgene in the Tg-rasH2 mouse model, a masked human genome was used for all classification where the mask territory was the exact sequence copy as integrated into Tg-rasH2.

Out of a total of 52,509,726 error-corrected paired-end reads across all 62 (1.2×10^−4^%) murine tissue samples, 50,910,333 were taxonomically classified as *Mus musculus,* 34 to *Rattus norvegicus albus,* 33 (6.3×10^−4^%) to *Homo sapiens,* and 0 to *Bos taurus* (0%). Exactly 84,865 (0.2%) paired-end reads were unclassified and 1,514,494 (2.8%) were from an ambiguous taxonomic origin. Only sequence data that could be positively identified as originating from the mouse genome was reserved for downstream analysis. Furthermore, every error-corrected paired-end read supporting a variant call in this cohort underwent manual review and BLAST+ alignment using the Blast nucleotide (nt) collection to confirm the true positive rate of taxonomic classification on this error-corrected dataset as being a perfect 100.000000%.

Tissue samples from vehicle control exposed mouse ID 9951 contained 29 paired-end reads from *Homo sapiens* and a tissue sample from the benzo[α]pyrene exposed mouse ID 9310 contained 28 paired-end reads from *Rattus norvegicus albus* suggesting that most contaminating events in both mouse cohorts were punctuated and private to just a few samples. The mean per-nucleotide mutant frequency for mouse 9951 is 1.2× 10^−7^ and if contaminating reads were not removed, the mean per-nucleotide mutant frequency would have risen to a rate equivalent, or greater than, the mutant frequencies detected in the positive control samples.

## SUPPLEMENTARY FIGURES

**Figure S1.**
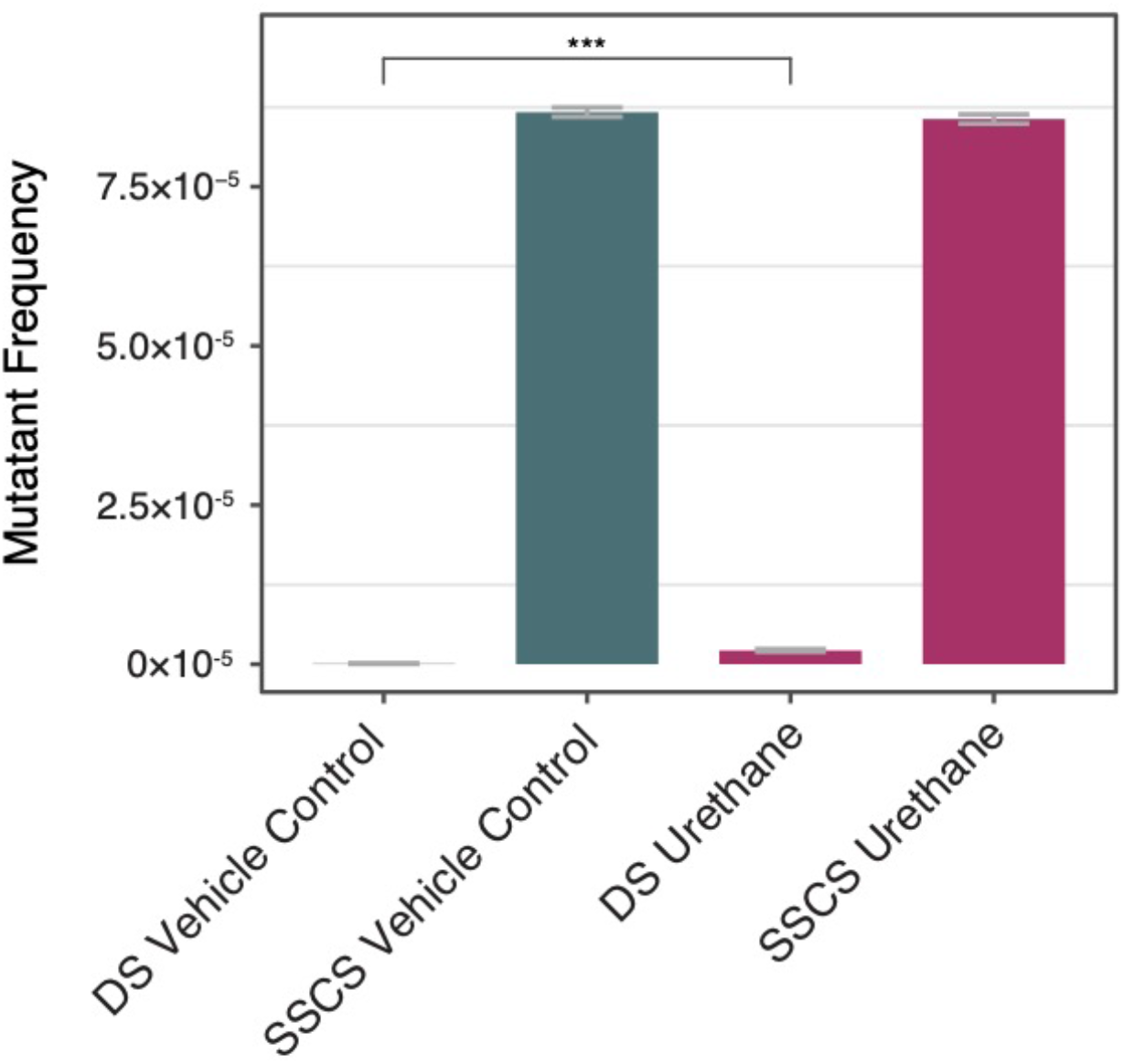
MF comparison in a mutagen exposed sample with and without duplex consensus level error-correction. Alternative forms of error-corrected next generation sequencing (ecNGS) may perform the error-correction on single-strands without resolving a complete duplex consensus. These single-strand error-correction forms of ecNGS are not sensitive enough for resolving small effect sizes in mutant frequency induction from experiments like those in the TGR assays. To illustrate this, we performed singlestrand error-correction data using Duplex Sequencing Adapters on two Tg-rasH2 mouse lung samples, one treated with urethane and one treated with the vehicle control. The per-nucleotide mutant frequencies for the vehicle control and urethane-exposed samples are 8.2×10^−8^ and 2.15×10^−6^ using Duplex Sequencing. When measuring the same metric using only single-strand consensus sequencing (SSCS), the two mutant frequencies rise to 8.6×10^−5^ and 8.6×10^−5^, respectively. The difference between the mutant frequencies of the exposed and control tissues using Duplex Sequencing are different with a p-value less than 2.2×10^−16^. This is in contrast to the single-strand error-correction measurements of mutant frequency which are not significant (p-value 0.98). Both statistical tests were performed using the Fisher’s exact test for count data. Error bars reflect 95% confidence intervals.

**Figure S2.**
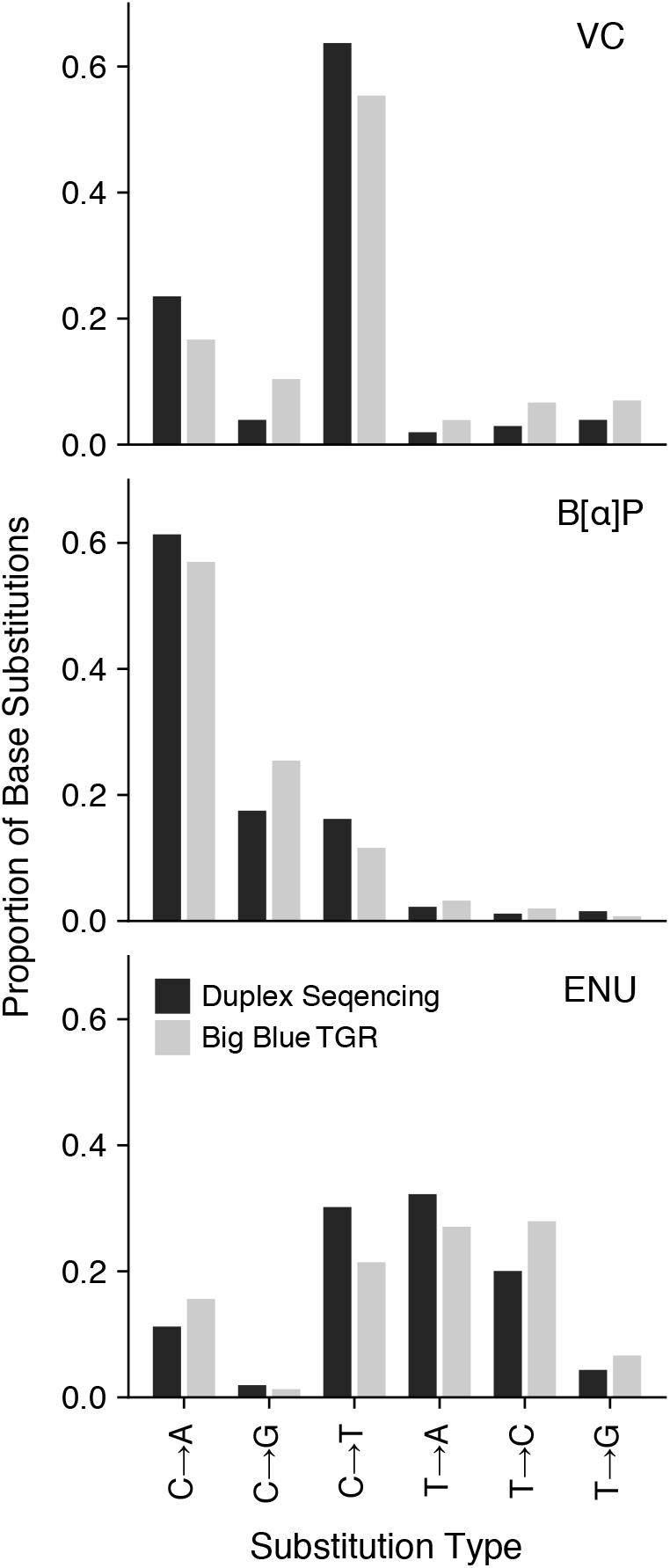
Mutation spectra observed by Duplex Sequencing of genomic DNA and individually sampled mutant plaques from the TGR assay are equivalent. The proportion of single nucleotide variants (SNV) within the *cII* gene are shown for individually picked mutant phage plaques produced from Big Blue rodent tissue and Duplex Sequencing of the *cII* transgene directly from gDNA. SNVs are designated with pyrimidine as the reference. The two methods yield the same spectrum for all treatment groups (p-values >0.999, chi-squared test). Proportions were calculated by dividing the total observations of SNVs by the total number of duplex bases within the *cII* interval and normalizing to one.

**Figure S3.**
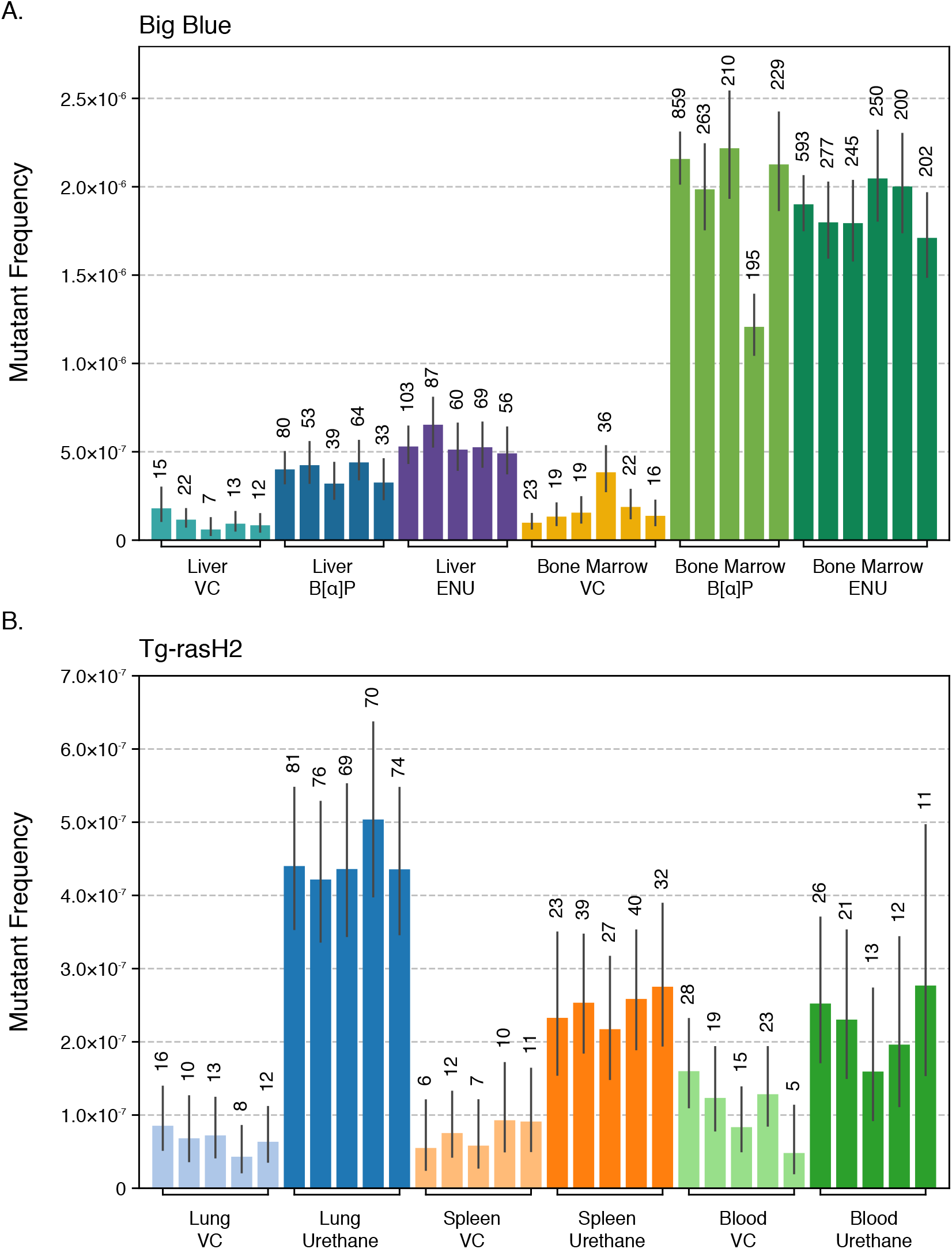
Per-sample mutant frequencies for all tissues and treatment groups. **A)** Big Blue cohort samples. **B)** Tg-rasH2 cohort samples. Mutant frequency was calculated as the total number of non-germline mutant duplex consensus base pairs divided by the total number of duplex consensus base pairs per sample. Error bars reflect 95% binomial confidence intervals. Integers above each bar represent the total number of mutant duplex consensus alleles observed per sample. VC, Vehicle Control.

**Figure S4.**
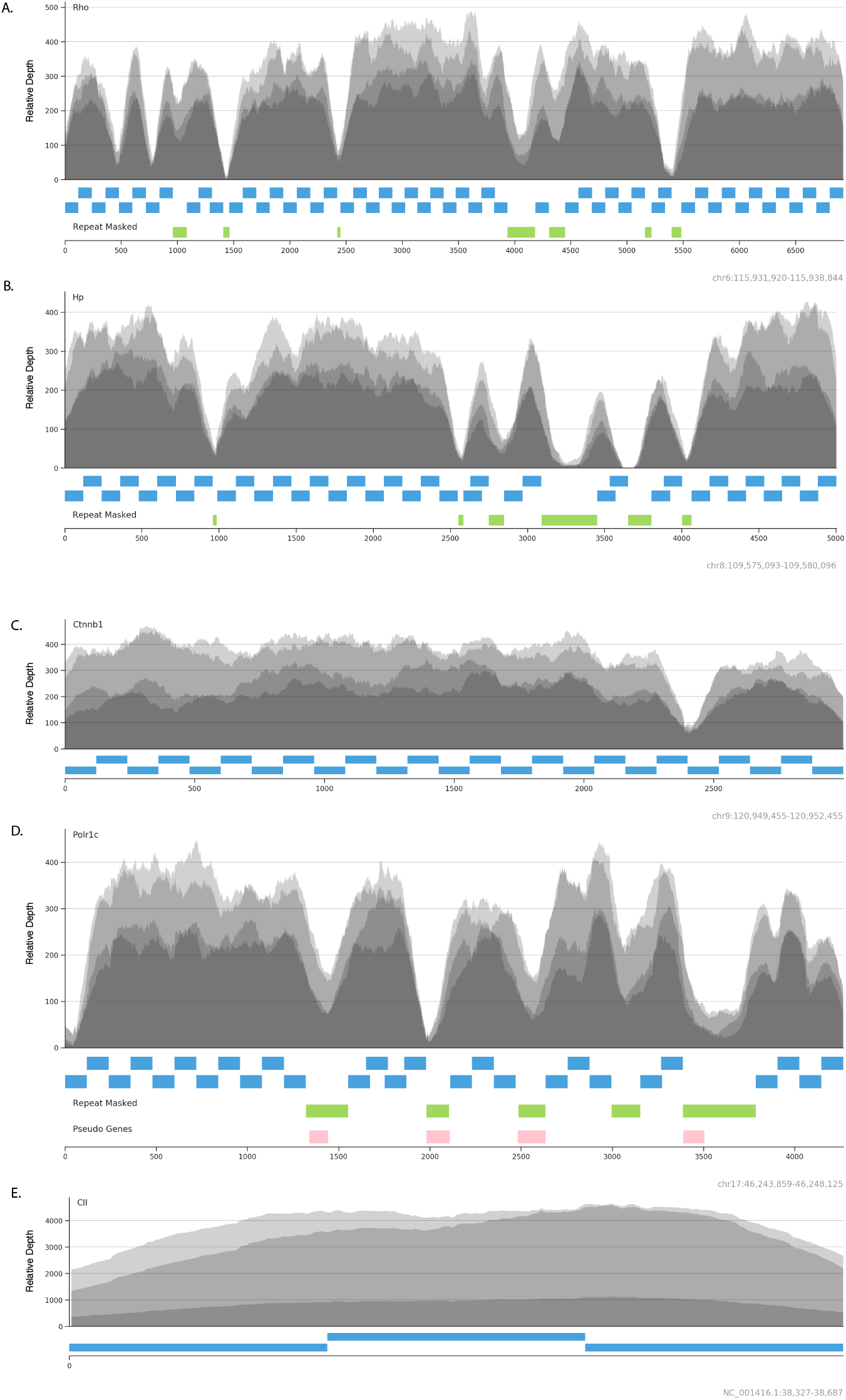
Consensus alignment data and probe design over endogenous and transgenic targets in the Big Blue mouse. Hybrid selection targets were carefully designed to abut no closer than 10 base pairs from a repeat masked (green) or pseudogene (pink) intervals. Individual baits are colored as blue intervals underlying the read coverage track. The four coverage tracks shown in all panels are from four randomly selected library preparations to illustrate the relatedness of coverage profile and bait layout. **A)** Example coverage and panel design over *Rho,* **B)** *Hp,* **C)** *Ctnnb1,* **D)** *Polr1c* and **E)** the Big Blue mouse *cII* transgene.

**Figure S5.**
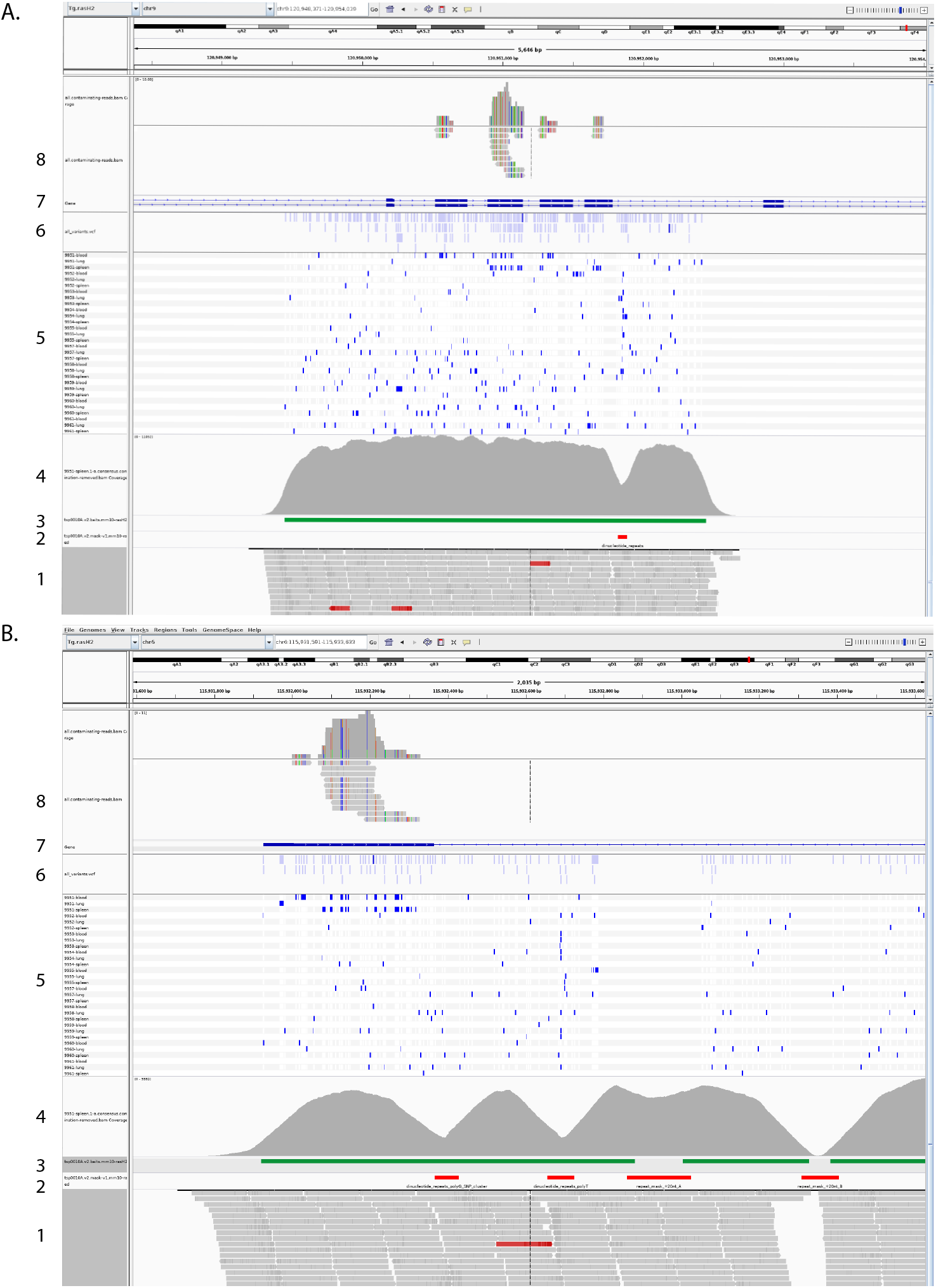
Ultra-rare contamination is easily detected by Duplex Sequencing but can be filtered from alignment data. Contamination of homologous species DNA drastically increases observed apparent mutations and confounds experimental frequency results in absence of identification and removal. **A**) A genomic view of the *Ctnnb1* gene in the transgenic mouse Tg-rasH2 (mm10). Tracks from bottom to top (1) consensus paired-end read alignments (2) repeat-masked regions (3) baited region (4) density of consensus paired-end read alignments (5) unfiltered variant calls for all samples (6) silhouette of unfiltered variant calls for all samples (7) transcript diagram of *Ctnnb1* (8) reads identified as contamination from all samples. **B**) A genomic view of the *Rho* gene in the transgenic mouse Tg-rasH2 (mm10). Tracks from bottom to top (1) consensus paired-end read alignments (2) repeat-masked regions (3) baited region (4) density of consensus paired-end read alignments (5) unfiltered variant calls for all samples (6) silhouette of unfiltered variant calls for all samples (7) transcript diagram of *Rho* (8) reads identified as contamination from all samples

## SUPPLEMENTARY TABLES

**Table S1:** Summary of *cII* mutant, and total, phage counts from Big Blue mouse samples assayed via the transgenic rodent assay.

| Treatment | Tissue | Big Blue Mouse ID | No. of Phage Screened | No. of Mutant Phage | Mutant Frequency ($\times 10^{-6}$) |
| --- | --- | --- | --- | --- | --- |
| Vehicle Control | Liver | 9301 | 795,000 | 32 | 40.3 |
|  |  | 9302 | 479,000 | 24 | 50.1 |
|  |  | 9303 | 851,667 | 36 | 42.3 |
|  |  | 9304 | 1,005,000 | 37 | 36.8 |
|  |  | 9305 | 1,090,000 | 57 | 52.3 |
|  |  | 9306 | 1,878,333 | 75 | 39.9 |
|  | Bone Marrow | 9301 | 1,056,667 | 43 | 40.7 |
|  |  | 9302 | 883,333 | 15 | 17.0 |
|  |  | 9303 | 945,000 | 27 | 28.6 |
|  |  | 9304 | 814,667 | 30 | 36.8 |
|  |  | 9305 | 452,333 | 22 | 48.6 |
|  |  | 9306 | 1,122,333 | 64 | 57.0 |
| Benzo[$\alpha$]pyrene | Liver | 9307 | 1061,667 | 183 | 172.4 |
|  |  | 9308 | 533,333 | 140 | 262.5 |
|  |  | 9309 | 1,048,333 | 212 | 202.2 |
|  |  | 9310 | 73,8333 | 159 | 215.3 |
|  |  | 9311 | 898,333 | 161 | 179.2 |
|  | Bone Marrow | 9307 | 461,667 | 359 | 777.6 |
|  |  | 9308 | 636,667 | 438 | 688.0 |
|  |  | 9309 | 982,167 | 563 | 573.2 |
|  |  | 9310 | 480,000 | 295 | 614.6 |
|  |  | 9311 | 486,667 | 356 | 731.5 |
| N-ethyl-N-nitrosourea | Liver | 9313 | 603,333 | 129 | 213.8 |
|  |  | 9314 | 268,000 | 62 | 231.3 |
|  |  | 9315 | 1,091,667 | 211 | 193.3 |
|  |  | 9316 | 926,667 | 128 | 138.1 |
|  |  | 9318 | 764,333 | 121 | 158.3 |
|  | Bone Marrow | 9313 | 1,310,000 | 413 | 315.3 |
|  |  | 9314 | 1,443,333 | 509 | 352.7 |
|  |  | 9315 | 1,081,667 | 336 | 310.6 |
|  |  | 9316 | 1,120,000 | 501 | 447.3 |
|  |  | 9317 | 555,333 | 279 | 502.4 |
|  |  | 9318 | 966,667 | 486 | 502.8 |

**Table S2.**
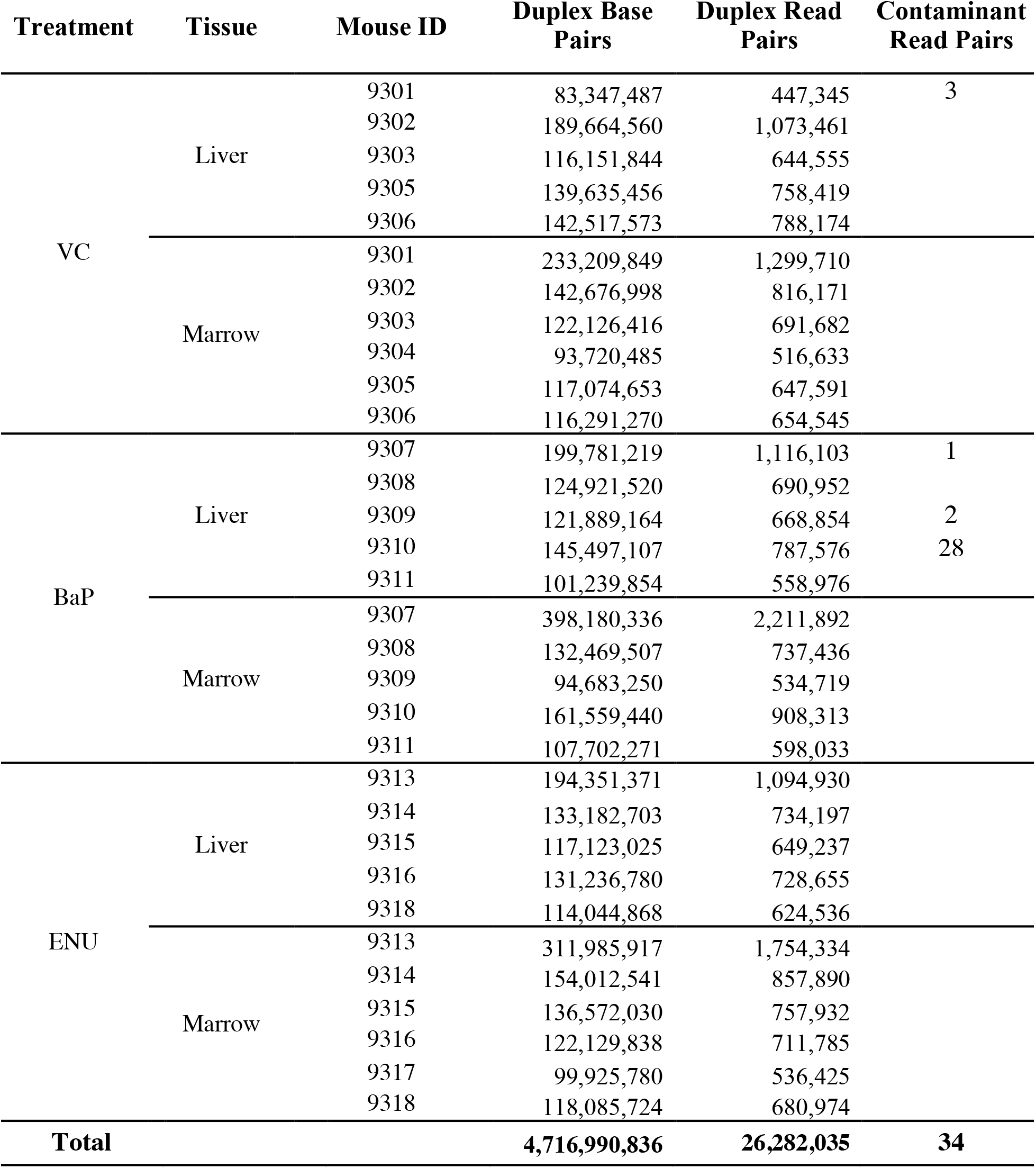
Sequencing summary of the Big Blue mouse samples including consensus duplex bases and read pairs assayed. Almost 5 billion duplex bases were generated from 26 million duplex consensus read pairs. Only 34 read pairs were positively assigned to either the *Rattus norvegicus albus* or *Homo sapiens* species and were removed prior to variant calling.

**Table S3:**
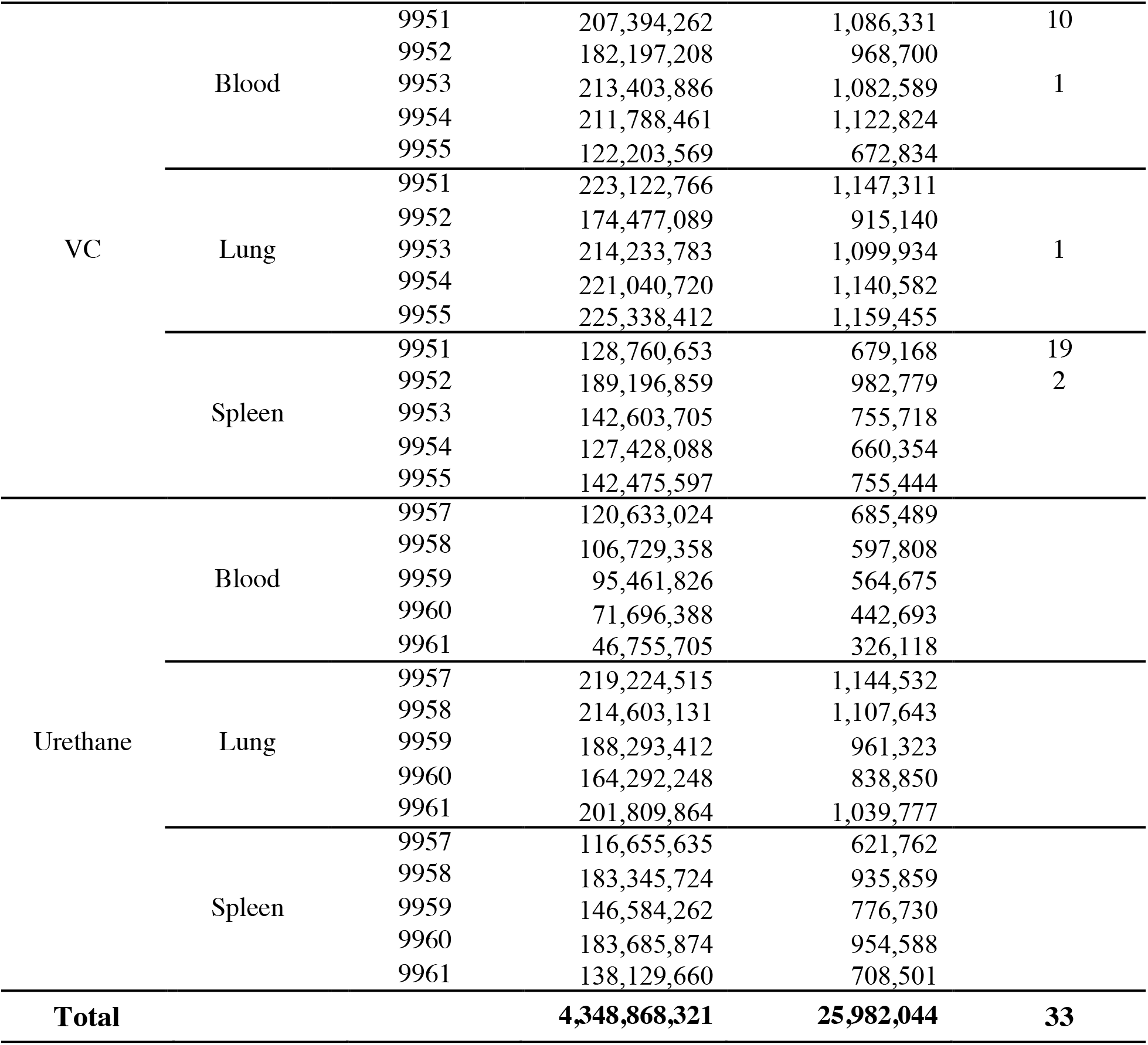
Sequencing summary of the Tg-rasH2 mouse samples including consensus duplex bases and read pairs assayed. Similar to the experimental design for sequencing of the Big Blue mouse samples, nearly 5 billion duplex base pairs were generated from 26 million duplex consensus read pairs from the Tg-rasH2 sample set. From these samples, 33 contaminating read pairs were detected from both *Rattus norvegicus albus* and *Homo sapiens* species. These reads were removed prior to downstream analysis.

**Table S4:**
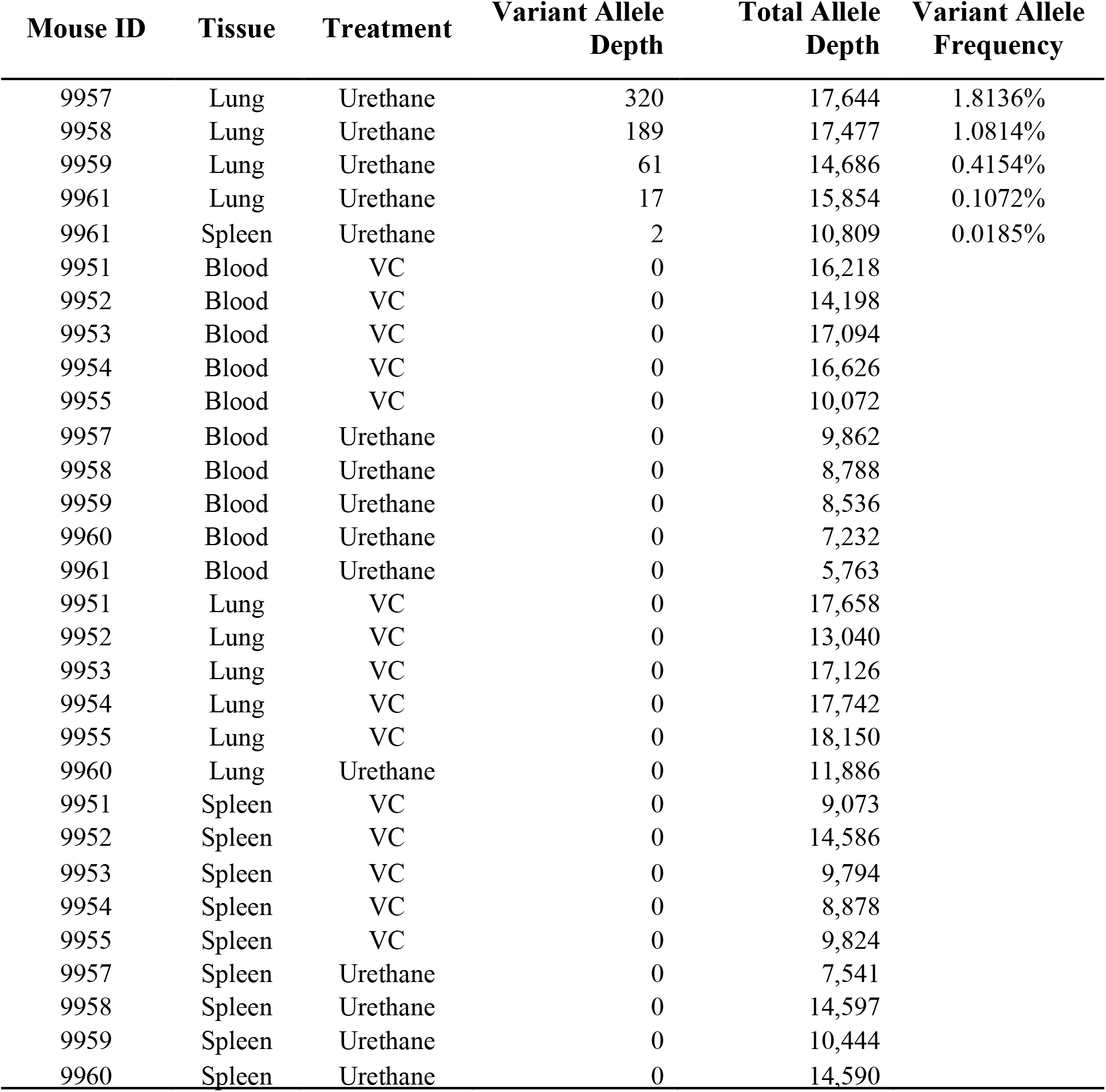
Early neoplastic evolution is detected with Duplex Sequencing in the cancer-predisposed mouse Tg-rasH2. The variant allele counts of A·T→T·A mutations at codon 61 in the human *HRAS* transgene in the Tg-rasH2 mouse model. The variant allele counts observed at this locus are those of A·T→T·A in the context CTG for urethane exposed tissues. All but one urethane exposed lung tissue harbors a variant at significant clonality. A single urethane exposed splenic sample has a small clone of two counts (0.018%) at this locus.

## SUPPLEMENTARY DATA FILES

**Database S1**. Tabular text file of all variant calls for the Big Blue samples in MUT format.

**Database S2**. Tabular text file of all variant calls for the Tg-rasH2 samples in MUT format.

